# Nance-Horan Syndrome-like 1 interacts with endophilin A2 and Ena/VASP proteins to promote fast endophilin-mediated endocytosis

**DOI:** 10.1101/2024.10.23.619882

**Authors:** Jonathan F. W. Cope, Ah-Lai Law, Samina Juma, Hayley J. Sharpe, Matthias Krause

## Abstract

Endocytosis, crucial for various physiological processes, facilitates receptor and extracellular material uptake. Fast endophilin-mediated endocytosis (FEME), driven by endophilin A2 (EndoA2), enables clathrin-independent, ligand-induced receptor uptake at the leading-edge of cells. Whilst F-actin polymerisation is essential for FEME, how actin dynamics are regulated to mediate FEME is unknown. NHSL1, a Nance-Horan Syndrome protein family member, localises to leading-edges of cells, where it regulates migration, and to vesicular puncta, where its function is undetermined. Here, we show that NHSL1 and its uncharacterised family member NHSL2 co-localise and engage in direct, multivalent interactions with EndoA2. NHSL1 also binds Ena/VASP proteins, a family of actin elongators. NHSL1 promotes FEME and its interactions with EndoA2 and Ena/VASP proteins are required for this function. NHSL1 does not control dynamin recruitment but enhances actin polymerisation at FEME sites. Thus, it may cooperate with EndoA2 and Ena/VASP proteins to control membrane invagination and actin polymerisation, thereby mediating FEME.

## Introduction

Endocytosis, a process by which cell surface molecules, bound cargos, and extracellular matter are internalised by the cell through the formation of membrane bound carriers, is crucial for embryonic development and tissue homeostasis. Whilst several distinct endocytic pathways have been identified, clathrin-dependent endocytosis is the most extensively studied and most widely utilised under constitutive conditions^1,2^. Nonetheless, multiple clathrin-independent endocytosis pathways are less well characterised, each occurring under specific conditions to uniquely contribute to cellular function^3,4^.

Fast endophilin-mediated endocytosis (FEME) is a clathrin-independent endocytosis pathway, which occurs at the leading edge of migrating cells and is triggered by stimulation of certain receptors, including the Epidermal growth factor (EGF) and beta1-adrenergic receptors, by their cognate ligands^5–7^. This pathway depends on membrane curvature induced by the endophilin A1 and A2 (subsequently referred to as EndoA) proteins^5^. This subfamily consists of endophilin A1 (EndoA1, also known as SH3GL2), endophilin A2 (EndoA2, also known as SH3GL1) and endophilin A3 (EndoA3, also known as SH3GL3), of which EndoA2 is the most ubiquitously expressed^8,9^. The EndoA proteins each harbour a BAR domain, which facilitates both their dimerisation and ability to bind to lipid bilayers and induce their curvature^10,11^. In addition, EndoA possesses a Src-homology 3 (SH3) domain which mediates its recruitment and clustering at FEME priming patches, cargo engagement and recruitment of dynamin and regulators of actin polymerisation^5,7,12–14^. The SH3 domain of EndoA exhibits the capability of binding to both canonical and non-canonical proline-rich SH3 domain-binding motifs (PRMs)^12,15–19^.

FEME requires a specialised membrane priming step, whereby a complex is assembled to cluster EndoA at specific plasma membrane sites called FEME priming patches^6,20^. At the leading edge of migrating cells, active CDC42 recruits the BAR domain containing proteins FBP17 and CIP4, which in turn recruit SHIP1, SHIP2 and Lamellipodin (Lpd, also known as RAPH1)^5,6^. Lpd contains ten proline-rich binding sites for the SH3 domain of EndoA2, which facilitate the recruitment and clustering of EndoA at sites of FEME priming^5,12^. In the absence of receptor stimulation, these priming patches disassemble, reforming elsewhere on the plasma membrane^6^. However, stimulation of local receptors drives their recruitment into FEME priming patches^5^. This triggers EndoA-induced membrane tubulation and local actin polymerisation which generates the force required for invagination, whilst dynamin- and friction-mediated scission facilitates FEME carrier budding^5,7,20–22^.

Whilst certain FEME players and their contribution to FEME are emerging, the detailed mechanism of FEME is still elusive. Furthermore, actin polymerisation has been shown to be essential for FEME^5^, but its regulation at sites of FEME and its precise role in membrane invagination and scission have yet to be studied. We recently showed that the actin regulators, VASP (vasodilator-stimulated phosphoprotein) and Mena (mammalian enabled) co-localise with EndoA2 at FEME priming patches^13^. VASP and Mena, alongside EVL (Ena/VASP-like), form the Ena/VASP protein family of actin elongators. These proteins have a conserved domain structure consisting of an EVH1 and EVH2 (Ena/VASP homology 1/2) domain connected by a central proline rich region (PRR)^23^. The EVH1 domain mediates their subcellular recruitment by binding to proline-rich peptides composed of a phenylalanine followed by four prolines and flanked by acidic amino acids (FP4 motifs) in other proteins such as Lpd^24–26^. We previously reported that the Ena/VASP proteins interact with Lpd to regulate both cell migration and the actin-dependent endocytosis of EGFR^12,23,26,27^. Furthermore, VASP, Lpd and EndoA form biomolecular condensates *in vitro*, with EndoA antagonising Lpd- and VASP-mediated actin polymerisation and bundling within these condensates^13^. These findings suggest that Ena/VASP proteins may contribute to the regulation of actin polymerisation during FEME.

Nance Horan Syndrome (NHS), NHS-like 1 (NHSL1), and NHS-like 2 (NHSL2) form the NHS protein family, a poorly characterised family of actin regulators^28–30^. Here we show that NHSL1 and NHSL2 both contain multiple proline-rich binding sites that facilitate a direct interaction of each protein with the SH3 domain of EndoA2. Furthermore, NHSL1 harbours two FP4 motifs which interact with the EVH1 domain of Ena/VASP proteins. We reveal that depletion of NHSL1 impairs FEME carrier formation, and that restoring NHSL1 expression with mutants impaired in binding to EndoA2 and Ena/VASP proteins cannot rescue these defects. Importantly, loss of NHSL1 does not affect dynamin recruitment but compromises F-actin polymerisation at FEME sites. Together these findings highlight a novel role for NHSL1 in FEME, which is dependent on its interactions with EndoA2 and Ena/VASP proteins and on its ability to control F-actin polymerisation.

## Results

### NHSL1 and NHSL2 co-localise with EndoA2 at vesicular puncta

We previously showed that NHSL1 negatively regulates cell migration through an interaction with the Scar/WAVE complex^29^. This interaction requires two proline-rich binding sites which mediate a direct interaction with the SH3 domain of Abi, a subunit of the Scar/WAVE complex^29^. Interestingly, NHSL1 localises not only to the edge of lamellipodia but also to vesicular puncta in migrating cells^29^, which suggests a role for NHSL1 in endocytosis or vesicle trafficking. To investigate whether this localisation is conserved amongst other members of the NHS protein family, we expressed mScarlet-tagged NHSL2 in B16-F1 cells plated on laminin. The B16-F1 line, a mouse melanoma cell line, was used in this context due to its ability to form large, flat lamellipodia^31^. We found that NHSL2, like NHSL1, localised to the very edge of the lamellipodium as well as to three separate populations of vesicular puncta: those near the leading edge, a central band anterior to the nucleus, and those at the rear of the cell (Fig. S1A, Supplementary Movie 1).

To explore the function of NHSL1 and NHSL2 at vesicles, we aimed to identify known regulators of endocytosis or vesicle trafficking that colocalise with NHSL1 or NHSL2 at vesicular puncta. As some NHSL1- and NHSL2-positive vesicles form in close proximity to the leading edge, we speculated that these may represent nascent FEME carriers. EndoA co-localises with the actin regulator Lpd, another interactor of the Scar/WAVE complex^32^, at clathrin coated pits and FEME priming patches^5,12^. Therefore, to establish whether similar co-localisation could be observed with NHSL1 and NHSL2, we co-expressed mScarlet-tagged NHSL1 (Fig. 1A, S1C, Supplementary Movie 2, 4) or NHSL2 (Fig. 1B, S1D, Supplementary Movie 3, 5) with EGFP-tagged EndoA2, in B16-F1 or BSC1 cells. BSC1 cells are commonly used to study FEME as they exhibit FEME priming patches and FEME carriers under both resting conditions and in response to stimulation with serum or various ligands^5,7^. We observed that EndoA2 localises to both NHSL1-and NHSL2-positive vesicles.

**Figure 1.**
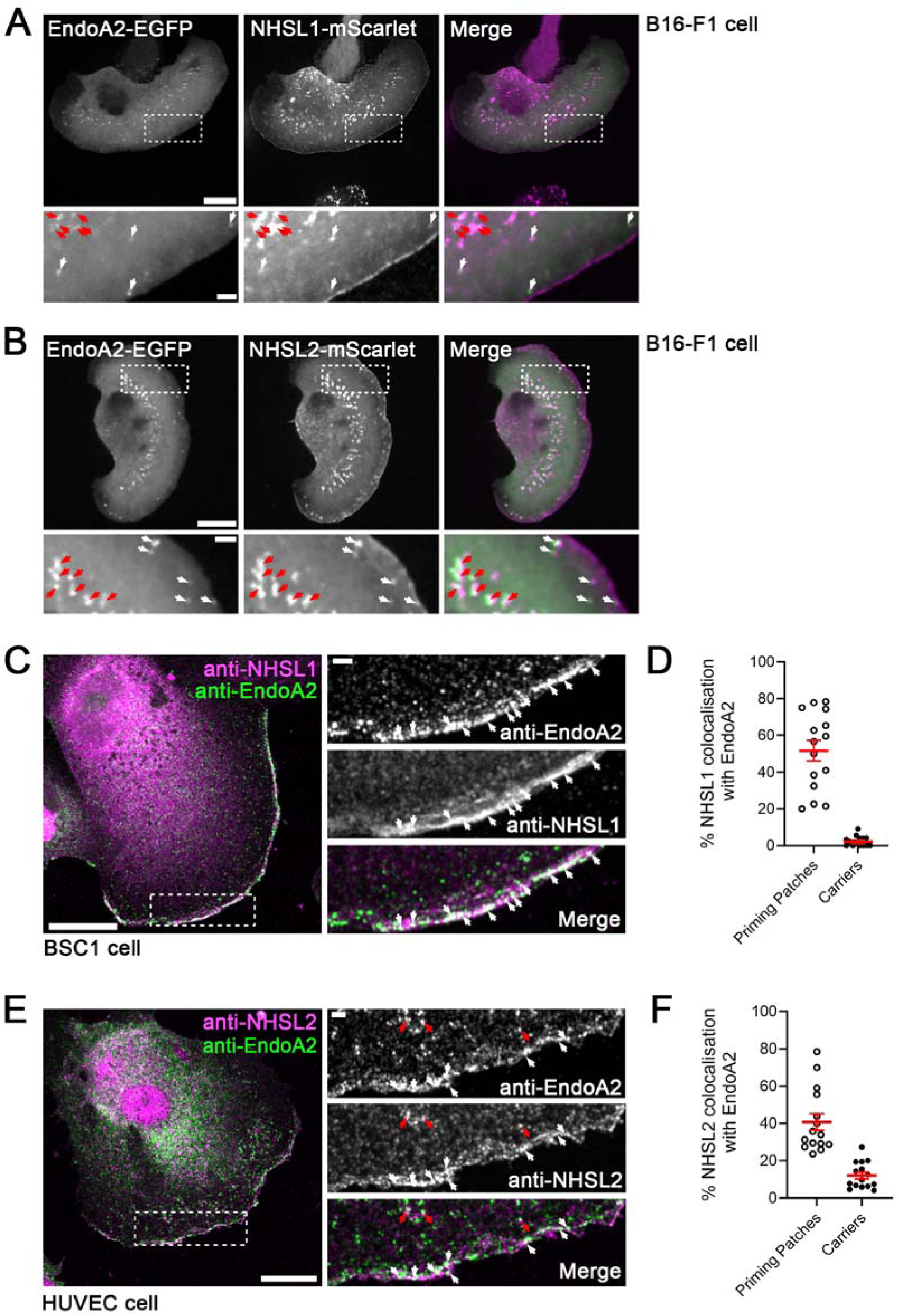
NHSL1 and NHSL2 co-localise with EndoA2. (**A** & **B**) B16-F1 cells were co-transfected with EGFP-tagged EndoA2 and mScarlet-tagged NHSL1 **(A, Supplementary Movie 2**) or NHSL2 (**B**, **Supplementary Movie 3**) and plated on laminin. Localisation of tagged proteins was imaged using live confocal microscopy. White and red arrows mark distinct populations of vesicular puncta. In merged stills, mScarlet-tagged NHSL1 and NHSL2 are represented in magenta, while EGFP-tagged EndoA2 is represented in green. Scale bars: 10 µm. Scale bar of insets: 2 µm. Images representative of three independent experiments. (**C**) Confocal images of a resting BSC1 cell, immuno-stained for endogenous EndoA2 and NHSL1 and imaged at a confocal plane placed in the middle (in Z) of the lamellipodium. White arrows mark FEME priming patches at which co-localisation can be observed. Scale bar: 20 µm. Scale bar of inset: 2 µm. Image representative of three independent experiments. (**D**) The percentage of FEME priming patches and carriers at which NHSL1 co-localised with EndoA2 was quantified before and after shifting the EndoA2 channel by 10 pixels in the X and Y directions (**Fig. S1E**). Graph shows normalised percentages calculated by subtracting the pixel shifted values from the unshifted values. Data represent means ± SEM with individual data points shown. Each data point represents one cell. N = 15 cells across 3 biological replicates. (**E**) Confocal image of a resting HUVEC cell, immuno-stained for endogenous EndoA2 and NHSL2 and imaged at a confocal plane placed in the middle (in Z) of the lamellipodium. White arrows mark FEME priming patches at which co-localisation can be observed. Red arrows mark FEME carriers at which co-localisation can be observed. Scale bar: 20µm. Scale bar of inset: 2 µm. Image representative of three independent experiments. (**F**) The percentage of FEME priming patches and carriers at which NHSL2 co-localised with EndoA2 was quantified before and after shifting the EndoA2 channel by 10 pixels in the X and Y directions (**Fig. S1F**). Graph shows normalised percentages calculated by subtracting the pixel shifted values from the unshifted values. Data represent means ± SEM with individual data points shown. Each data point represents one cell. N = 15 cells across 3 biological replicates.

We next asked whether endogenous NHSL1 or NHSL2 could be found at FEME priming patches and/or FEME carriers in cells. We first investigated endogenous protein levels of NHSL1, NHSL2 and EndoA2 in small panel of cell lines including BSC1, B16-F1 and human vascular endothelial cells (HUVEC) (Fig. S1B). HUVEC, like BSC1 cells, exhibit high FEME activity^7^. We found that NHSL1 is expressed at low levels in B16-F1 and BSC1 cells, whereas it is highly expressed in HUVEC. Furthermore, NHSL2 expression, whilst not detectable by western blot in B16-F1 and BSC1 cells, is highly expressed in HUVEC. Therefore, we decided to explore NHSL1 and NHSL2 localisation in BSC1 cells and HUVEC respectively.

For our co-localisation analysis, we defined FEME priming patches as EndoA2-positive assemblies (EPAs) at the very edge of lamellipodia and FEME carriers as EPAs at least 1 µm from the leading edge, as detected in a confocal plane placed at the middle (in Z) of the lamellipodium. We found that endogenous NHSL1 or NHSL2 colocalised with EndoA2 at ∼52% or 41% of FEME priming patches respectively, as well as at a small proportion (∼2% for NHSL1 or 12% for NHSL2) of FEME carriers (Fig. 1C-F controlled for background; S1E,F raw data).

### The SH3 domain of EndoA2 binds directly to multiple sites in NHSL1 and NHSL2

Having observed co-localisation of NHSL1 and NHSL2 with EndoA2, we next wanted to establish whether these proteins interact. We co-expressed Myc-tagged EndoA2 with two EGFP-tagged NHSL1 isoforms (A1 & F1) or NHSL2 in HEK293 cells. NHSL1-A1 and NHSL1-F1 differ only in their first exon, with NHSL1-A1 harbouring an exon one which contains a Scar/WAVE homology domain^30^. Using EGFP pulldown experiments, we found that Myc-EndoA2 associated with both EGFP-tagged NHSL1 isoforms, as well as with NHSL2 (Fig. 2A). We observed that NHSL1-F1 had a higher affinity for EndoA2 (Fig. 2A) and thus we focussed on this isoform in the remainder of this study (all further exogenously expressed NHSL1 refers specifically to the NHSL1-F1 isoform). Furthermore, we found that NHSL2 can interact with all three EndoA proteins (Fig. 2B).

**Figure 2.**
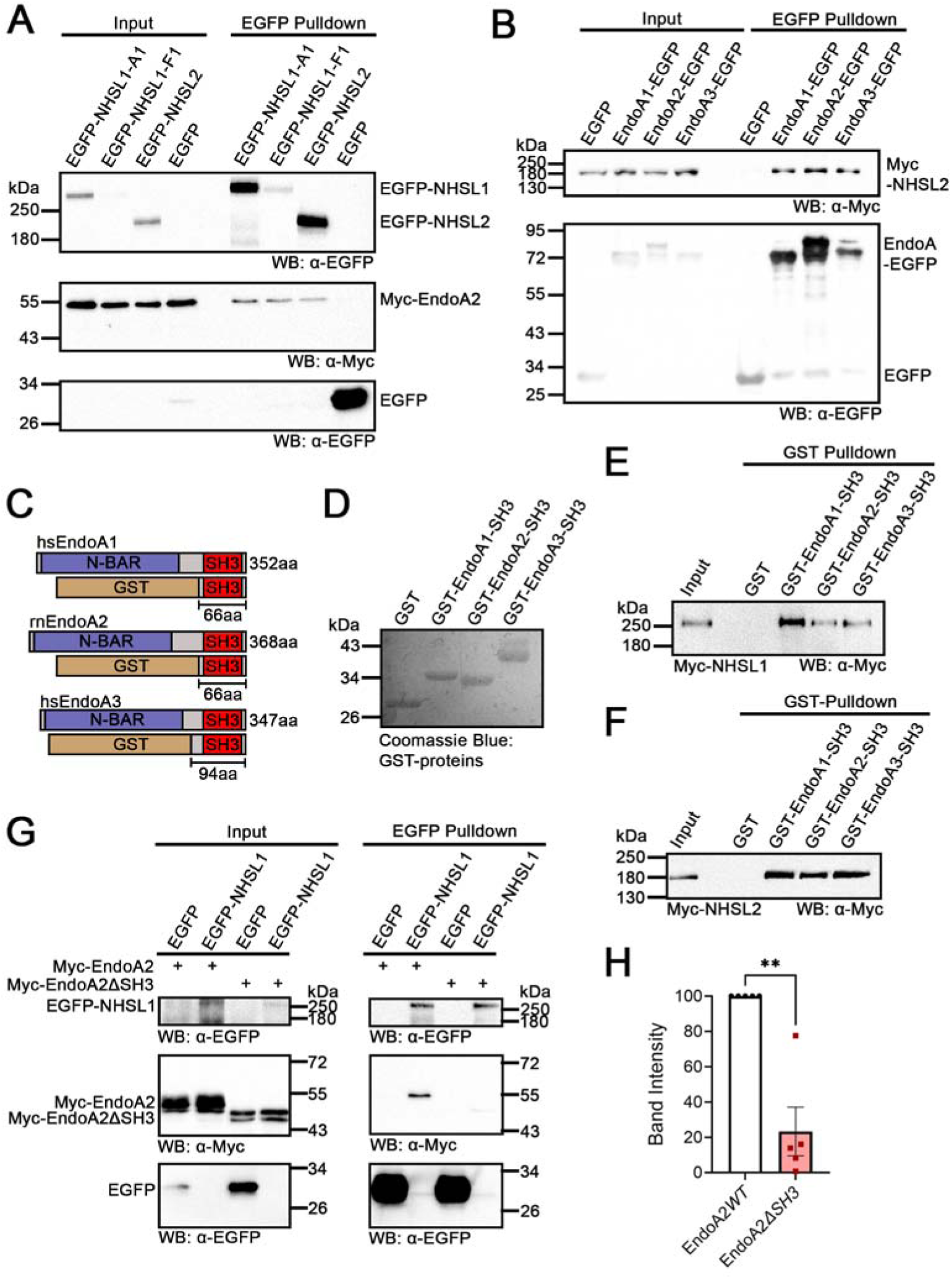
NHSL1 and NHSL2 interact with the SH3-domains of EndoA1-3. (**A**) Myc-EndoA2 was co-expressed with EGFP-NHSL1-A1, EGFP-NHSL1-F1, EGFP-NHSL2 or EGFP alone in HEK293 cells. EGFP-tagged proteins were pulled down from lysates using a nanobody against EGFP and blots were probed using antibodies against Myc and EGFP. Blot representative of three independent experiments. (**B**) Myc-NHSL2 was co-expressed with EndoA1-EGFP, EndoA2-EGFP, EndoA3-EGFP or EGFP alone in HEK293 cells. EGFP-tagged proteins were pulled down from lysates using a nanobody against EGFP and blots were probed using antibodies against Myc and EGFP. Blot representative of three independent experiments. (**C**) Schematic showing domain structure of EndoA1-3 and GST-fusion proteins used in **D**-**F** (**D**) GST-tagged SH3 domains of EndoA1 (GST-EndoA1-SH3), EndoA2 (GST-EndoA2-SH3), EndoA3 (GST EndoA3-SH3) as well as GST only (GST) were purified from *E. coli* using glutathione beads. Purified proteins were visualised with Coomassie blue stain. Note the higher molecular weight of GST-EndoA3-SH3 due to the inclusion of more amino acids directly upstream of the SH3 domain start site. (**E**-**F**) Myc-NHSL1 (**E**) or Myc-NHSL2 (**F**) were expressed in HEK293 cells. Purified GST-tagged constructs or GST only were used to pull down associated proteins from lysates and blots were probed using an antibody against Myc. Blots representative of three individual experiments. (**G**) EGFP-NHSL1 or EGFP were co-expressed with either Myc-tagged wild-type EndoA2 (Myc-EndoA2*WT*) or Myc-tagged EndoA2 with a deletion to its entire SH3 domain (Myc-EndoA2Δ*SH3*) in HEK293 cells. Proteins were pulled down from lysates using a nanobody against EGFP and blots were probed using antibodies against Myc and EGFP. Blot representative of five independent experiments. (**H**) Quantification of band intensity from **G**. EGFP pulldown bands from α-Myc panel were first normalised to the corresponding α-Myc input band to account for variability in expression and then normalised to the corresponding pulldown band from the α-EGFP panel. Each value was then normalised to that of EndoA2*WT* which was set to 100%. Results represent means ± SEM with individual data points from each replicate shown. N_=_5 independent biological experiments. Statistics performed using a student’s two-tailed t-test: **P = 0.0052.

Both NHSL1 and NHSL2 are largely unstructured and contain multiple proline-rich motifs which facilitate binding to proteins that contain SH3 domains (Tables S1, S2)^33^. Therefore, to determine whether the interaction of EndoA with NHSL1 and NHSL2 is mediated by its SH3 domain, we purified GST-tagged SH3 domains of EndoA1-3 from *E. coli* (Fig. 2C, D). We found that the SH3 domains of all three EndoA proteins were able to interact with Myc-tagged NHSL1 (Fig. 2E) and Myc-tagged NHSL2 (Fig. 2F). To investigate whether the SH3 domain is necessary for the interaction of EndoA with NHSL1, we generated a mutant EndoA2 construct which lacked the SH3 domain (Myc-EndoA2-ΔSH3). We found that in the absence of its SH3 domain, the interaction of EndoA2 with NHSL1 is significantly disrupted (Fig. 2G, H). Together, these data indicate that the SH3 domain of EndoA2 is sufficient for its interaction with NHSL1 and NHSL2 and is necessary for its interaction with NHSL1.

We next wanted to identify whether the SH3 domain of EndoA2 interacts directly with NHSL1. To achieve this, we purified MBP-tagged EndoA2-SH3 (Fig. S2A) and eight GST-tagged fragments spanning the length of NHSL1-F1 (Fig. S2B,C) for use in far-western blot experiments. Far-western blots are used to detect direct protein-protein interactions through the use of a protein probe (in this case MBP-EndoA2-SH3 or MBP) prior to using a primary and secondary antibody^34^. We noticed that NHSL1 fragments 2 and 3 show substantial degradation, but together they contain only one putative binding site, which spans their intersection (Fig. S2C). We found that the EndoA2 SH3 domain bound directly to multiple fragments of NHSL1, binding most strongly to fragments 4, 5, 6 and 8 and weakly to fragment 1 (Fig. 3A). Additionally, we observed absence of non-specific binding of MBP to any fragment or to GST (Fig. S2D; imaged at the same time with the same parameters). To identify the exact sites which mediate the interaction between EndoA2 and NHSL1, we identified 33 putative PRMs in human NHSL1 using the online webtool ScanSite (https://scansite4.mit.edu) (Table S1)^35^. We assessed binding of the SH3 domain of EndoA2 to an array of 12mer peptides containing each of the 33 PRMs. The EndoA2 SH3 domain successfully bound to 13 of the 33 motifs (Fig. 3B), including the previously characterised Abi binding sites^29^. However, both Abi binding sites contain 2 individual SH3-domain binding motifs in close proximity, giving 11 separate binding sites in total (Fig. 3C).

**Figure 3.**
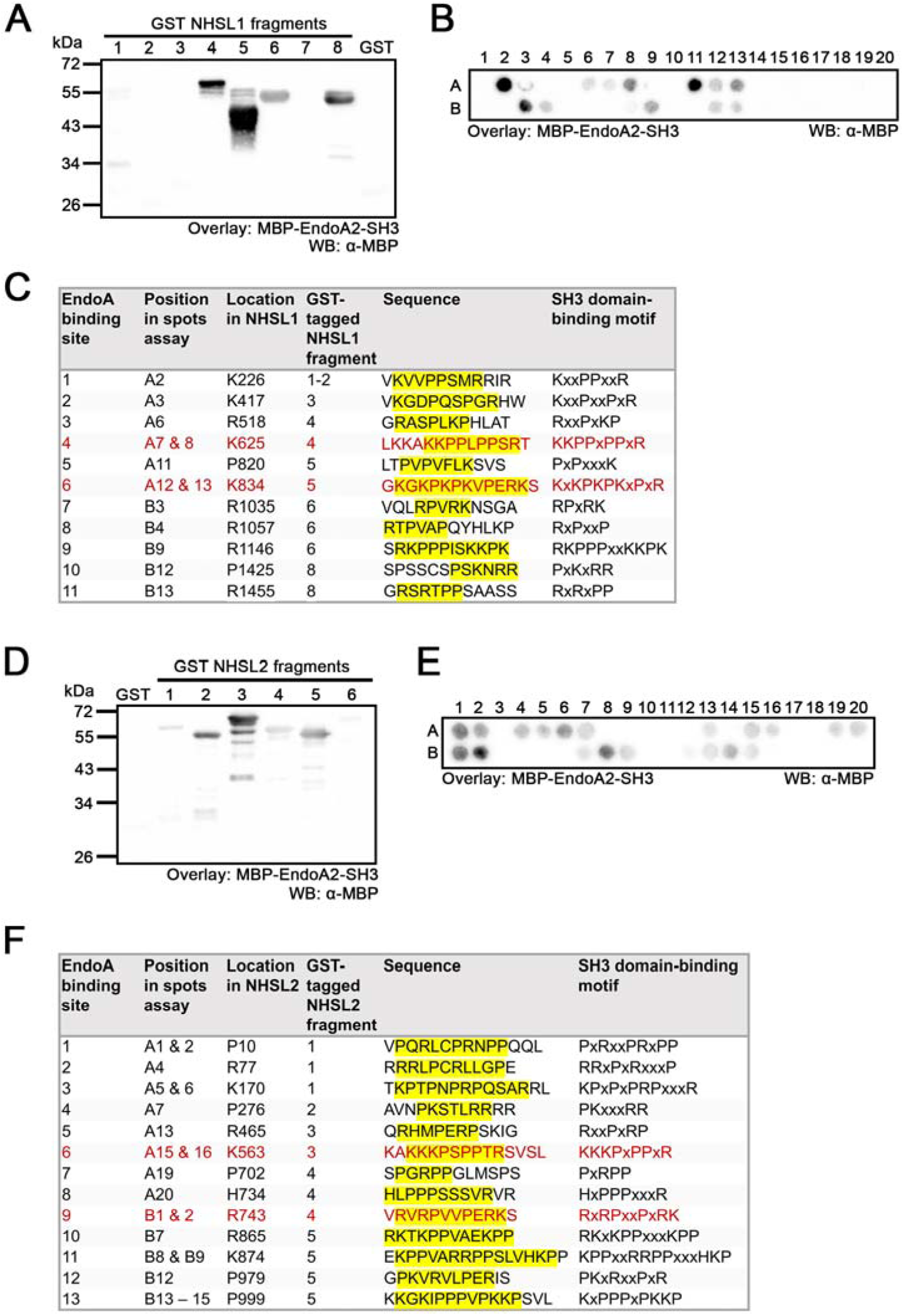
NHSL1 and NHSL2 harbour multiple binding sites for the SH3 domain of EndoA2. (A) Far western blot experiment using GST-tagged NHSL1-F1 fragments (see **Fig. S2B, C**) overlaid with MBP-tagged EndoA2 SH3 domain or MBP control (see **Fig. S2D,** exposed at the same time and with the same parameters) and probed using an antibody against MBP. Blots are representative of three individual experiments. (B) 33 putative SH3-domain binding motifs in NHSL1 (see **Table S1**) were synthesized as peptide spots on a membrane. The membrane was probed with purified MBP-tagged EndoA2 SH3 domain followed by an antibody against MBP. A1-A20 and B1-B13 represent a grid to easily identify individual peptides. (**C**) Table summarising positive hits from **B**. Multiple motifs that are in close enough proximity (< 6 aa) to one another to constitute one individual binding site have been grouped together. Previously characterised Abi binding sites^29^ are shown in red. (**D**) Far western blot experiment using GST-tagged NHSL2 fragments (see **Fig. S2E, F**) overlaid with MBP-tagged EndoA2 SH3 domain or MBP control (see **Fig. S2G,** exposed at the same time and using the same parameters) and probed using an antibody against MBP. Blots are representative of three individual experiments. (**E**) 38 putative SH3-domain binding motifs in NHSL2 (see **Table S2**) were synthesized as peptide spots on a membrane. The membrane was probed with purified MBP-tagged EndoA2 SH3 domain followed by an antibody against MBP. A1 – A20 and B1 – B13 represent a grid to easily identify individual peptides. (**F**) Table summarising positive hits from **E**. Multiple motifs that are in close enough proximity (<6 aa) to one another to constitute one individual binding site have been grouped together. Conserved Abi binding sites are shown in red.

To identify the sites to which the EndoA2 SH3 domain binds in NHSL2, we performed an identical set of experiments to those used to elucidate the 11 sites in NHSL1. We first purified six fragments spanning the length of NHSL2 (Fig. S2E,F). Using far western experiments, we found that the EndoA2 SH3 domain bound directly to multiple fragments of NHSL2, binding most strongly to fragments 3 and 5, weakly to fragments 1, 2 and 4 and minimally to fragment 6 (Fig. 2D). MBP bound weakly and non-specifically to fragments 2 and 3, although it did not bind to GST alone (Fig. S2G; imaged at the same time with the same parameters). Using ScanSite, we identified 38 putative PRMs in human NHSL2 (Table S2). The EndoA2 SH3 domain displayed successful binding to 20 of these PRMs, which again included the two conserved Abi binding sites (Fig. 3E). Of these 20 motifs, several occur in such close proximity to one another within the sequence of NHSL2, that they cannot be referred to as separate binding sites. In these cases, two or three motifs have been combined to form one individual SH3 domain binding site, giving 13 binding sites in total (Fig. 3F).

Taken together, these data suggest that EndoA2 interacts with NHSL1 and NHSL2, with its SH3 domain binding directly to multiple PRMs within each protein which may increase local concentration of EndoA2 BAR domains to induce membrane invagination.

### NHSL1 interacts with Ena/VASP proteins via two EVH1 domain-binding motifs

VASP interacts with EndoA1 and Lpd, with multivalent interactions between these proteins driving the formation of biomolecular condensates, through liquid-liquid phase separation (LLPS), *in vitro*^13^. Furthermore, we found that VASP and Mena both co-localise with EndoA2 at FEME priming patches^13^. Interestingly, analysis of the NHSL1 amino acid sequence revealed that it contains two putative FP4 motifs, which are known to interact with the EVH1 domain of Ena/VASP proteins^24,25^. Therefore, we speculated that NHSL1 may interact with Ena/VASP proteins to contribute to the regulation of actin polymerisation at sites of FEME.

To establish whether NHSL1 co-localises with Ena/VASP proteins in cells, we co-expressed EGFP-tagged NHSL1-F1 with mScarlet-tagged EVL, VASP and Mena in B16-F1 cells. We observed co-localisation of NHSL1-F1 with EVL (Fig. S3A) and VASP (Fig. S3B) at vesicular puncta and with all three Ena/VASP proteins at the edge of lamellipodia (Fig. S3A - C, Supplementary Movie 6 - 8). To test whether NHSL1 and Ena/VASP proteins can interact, we co-expressed EGFP-NHSL1-F1 with Myc-tagged VASP, Mena or EVL in HEK293 cells. We pulled-down EGFP-NHSL1-F1 from cell lysates and found that NHSL1 interacts with all three Ena/VASP proteins (Fig. 4A). We next performed endogenous co-immunoprecipitation experiments using an antibody against all NHSL1 isoforms and observed that endogenous NHSL1 associates with EVL (Fig. 4B).

**Figure 4.**
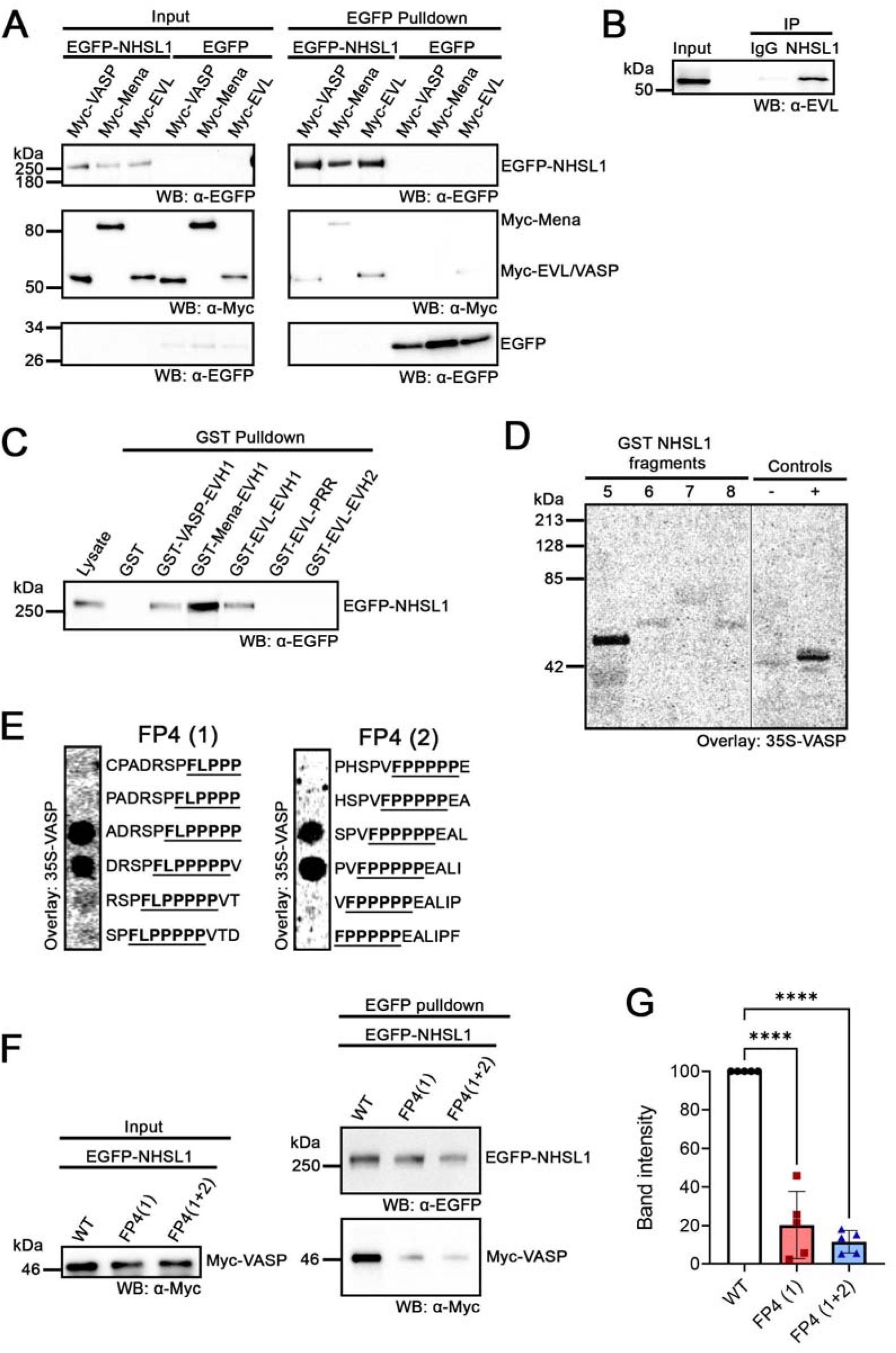
NHSL1 interacts with Ena/VASP proteins via two FP4 motifs. (**A**) Myc-VASP, Myc-Mena or Myc-EVL were co-expressed with EGFP-NHSL1-F1 or EGFP alone in HEK293 cells. Lysates were pulled down using a nanobody against EGFP and blots were probed using antibodies against Myc and EGFP. Blot representative of five independent experiments. (**B**) Endogenous NHSL1 was immunoprecipitated with NHSL1 polyclonal antiserum from B16-F1 lysates, blotted and probed using an antibody against EVL. Blot representative of 3 independent experiments. (**C**) EGFP-NHSL1-F1 was expressed in HEK293 cells. Purified GST-tagged constructs (Fig. S3) or GST only were used to pull down associated proteins from lysates and blots were probed using an antibody against EGFP. Blots representative of three individual experiments. (**D**) Far western blot experiment using four GST-tagged NHSL1 fragments (Fragment 5 - 8 in Fig. S2B,C) overlaid with ^35^S-labelled VASP. Bound VASP was detected using a Phosphorimager. A GST-fusion protein containing four FP4 motifs of the Listeria ActA protein served as a positive (+) control and GST alone as the negative (−) control. (**E**) The indicated peptides were synthesized as peptide spots on a membrane. The membrane was probed with ^35^S-labelled VASP and imaged using a Phosphorimager. The location of the FP4 motif is underlined in each peptide. (**F**) Myc-tagged VASP was co-expressed with EGFP-tagged wild-type NHSL1-F1, EGFP-tagged NHSL1-F1 with the first FP4 motif mutated (FP4 (1), FLPPPPP>ALAAPPP), or EGFP-tagged NHSL1-F1 with both FP4 motifs mutated (FP4 (1+2), FP4-1: FLPPPPP>ALAAPPP, FP4-2: VFPPPPP>AAAAPPP). Lysates were pulled down using a nanobody against EGFP and blots were probed using antibodies against Myc and EGFP. Blot representative of five independent experiments. (**G**) Quantification of band intensity from **F**. Bands from α-Myc panel were normalised to the corresponding band from α-EGFP panel. Each value was then normalised to that of wild-type NHSL1-F1 which was set to 100%. Results represent means ± SEM with individual data points from each replicate shown. N_=_5 independent biological experiments. Statistics performed using one-way ANOVA with Dunnett’s multiple comparisons test: ****P < 0.0001.

To investigate whether the EVH1 domain of Ena/VASP proteins mediates their interaction with NHSL1, we purified GST-tagged EVH1 domains from EVL, VASP, and Mena, as well as the PRR and EVH2 domains of EVL, all from *E. coli* (Fig. S4). Neither the EVL PRR nor EVH2 domain pulled down NHSL1-F1 from HEK293 cell lysates, indicating that these domains are not needed for the interaction of Ena/VASP proteins with NHSL1 (Fig. 4C). In contrast, the EVH1 domains of all three Ena/VASP proteins bound NHSL1-F1, suggesting that Ena/VASP proteins interact with NHSL1 via their EVH1 domain (Fig. 4C). We next probed the four C-terminal NHSL1 fragments (Fragment 5 - 8 in Fig. S2B,C) for their ability to interact directly with ^35^S-labeled, *in vitro* translated VASP. A GST-fusion protein containing four FP4 motifs of the *Listeria monocytogenes* ActA protein, a well-characterised Ena/VASP ligand^24^, served as a positive control and GST alone as the negative control. We found that ^35^S-labeled VASP bound to NHSL1 fragment 5, which contains both of the FP4 motifs found in NHSL1, indicating that they interact (Fig. 4D). To further map the Ena/VASP binding sites within this fragment of NHSL1, we used a peptide array with overlapping peptides covering both FP4 motifs and adjacent amino acids and overlaid it with ^35^S-labeled VASP. This indicates that the N-terminal aspartic acid residue at position −4 before the first FP4 motif and at least 5 prolines within this motif are essential for VASP binding, whilst binding to the second FP4 motif requires at least two N-terminal amino acids before the phenylalanine and a C-terminal leucine at position F+8 (Fig. 4E). This is consistent with previous studies reporting that acidic amino acids flanking FP4 motifs are essential for specificity of the interaction with the EVH1 domain of Ena/VASP proteins^24,25^.

To further validate the two mapped EVH1 binding sites, phenylalanine and selected prolines in each FP4 motif were mutated to alanine in the full-length cDNA of NHSL1-F1 (FP4-1: FLPPPPP>ALAAPPP; FP4-2: VFPPPPP>AAAAPPP). We co-expressed Myc-tagged VASP with EGFP-tagged wild-type NHSL1-F1 (EGFP-NHSL1*WT*), EGFP-tagged NHSL1-F1 with a mutation in the first FP4 site (EGFP-NHSL1*FP4(1)*), or EGFP-tagged NHSL1-F1 with mutations in both FP4 sites (EGFP-NHSL1*FP4(1+2*)), in HEK293 cells. We found that mutating either the first FP4 site, or both FP4 sites significantly disrupted the interaction of VASP with NHSL1 (Fig. 4F, G). Taken together, these results highlight that NHSL1 harbours two FP4 motifs, which mediate its direct interaction with the EVH1 domains of all three Ena/VASP proteins.

### Mutating the EndoA2 and Ena/VASP binding sites disrupts the interaction between EndoA2 and NHSL1

Having identified 11 sites in NHSL1 that bind to the SH3 domain of EndoA2, we next assessed whether these sites mediate the interaction between EndoA2 and full length NHSL1. Each SH3-domain binding site was mutated by substituting every proline in the PRM with an alanine to generate a mutant NHSL1-F1 construct in which all 11 sites were mutated (NHSL1*EndoAbi*), as well as a second construct in which only the 9 EndoA2 sites that do not bind Abi were mutated (NHSL1*Endo*, Fig. 5A). This second construct was designed with future functional assays in mind, so that we could be confident that any observed phenotype was not caused by impaired binding of NHSL1 to the Scar/WAVE complex via Abi. As NHSL1 is largely unstructured we did not anticipate that introducing these mutations would significantly affect its global properties. Having generated both mutant constructs, we performed far western experiments to assess whether these mutations impaired direct binding of the EndoA2 SH3 domain to NHSL1. For these experiments, we included a third mutant with mutations exclusively at the two Abi binding sites (NHSL1*Abi*, Fig. 5A), to assess the individual contribution of these two sites to the interaction between NHSL1 and EndoA2. We found that binding of the EndoA2 SH3 domain to both NHSL1*EndoAbi* and NHSL1*Abi* was significantly impaired (Fig. 5B, C). In contrast, binding to NHSL1*Endo* was not significantly different from that of wild-type NHSL1 (Fig. 5B, C). This finding suggests that the two sites to which both EndoA2 and Abi can bind are particularly important for the direct interaction between the SH3 domain of EndoA2 and NHSL1.

**Figure 5.**
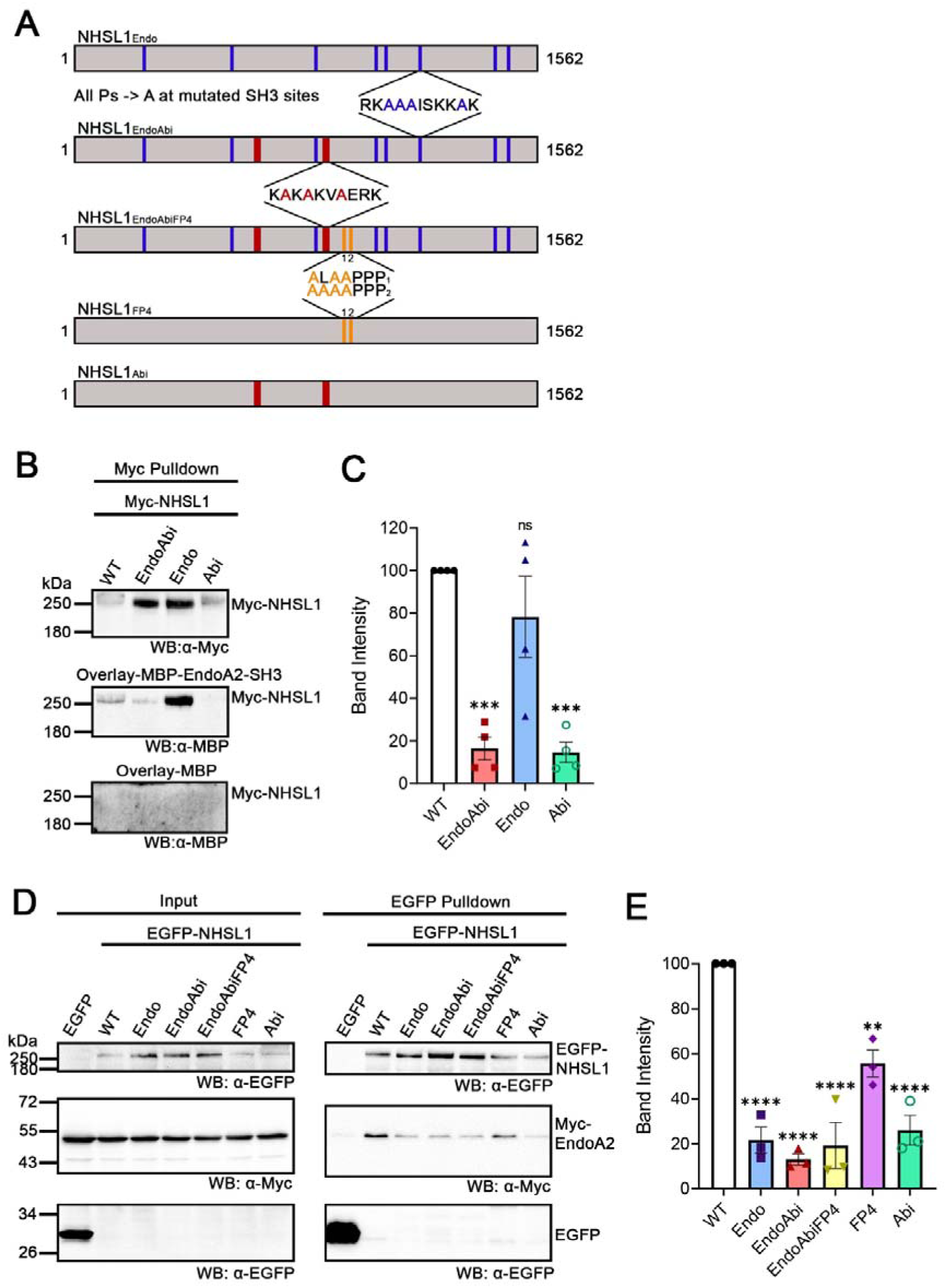
Mutating the EndoA2 and Ena/VASP binding sites disrupts the interaction of NHSL1 with EndoA2. **(A)** Schematic summarising the different EndoA2 and Ena/VASP binding mutants. The location of the mutated sites in full length NHSL1 are indicated. At each EndoA2 binding site, every proline was mutated to an alanine. Specific amino acid substitutions are shown for each FP4 site: FP4-1: FLPPPPP>ALAAPPP; FP4-2: VFPPPPP>AAAAPPP. (**B**) Far-western blot experiment using Myc-tagged wild-type (WT) or mutant NHSL1-F1 purified from HEK293 cell lysates using a nanobody against Myc. Blots were overlaid with MBP-tagged EndoA2 SH3 domain (middle panel) or MBP control (bottom panel) and probed using an antibody against MBP. Relative amount of purified WT or mutant NHSL1-F1 was visualised using an antibody against Myc (top panel). Blots are representative of four independent experiments. (**C**) Quantification of band intensity from **B**. Bands from Overlay-MBP-EndoA2-SH3 panel were normalised to the corresponding band from α-Myc panel. Each value was then normalised to that of wild-type NHSL1-F1 which was set to 100%. Results represent means ± SEM with individual data points from each replicate shown. N_=_4 independent biological experiments. Statistics performed using one-way ANOVA with Dunnett’s multiple comparisons test: *** P = 0.0002, ns P = 0.336. (**D**) Myc-EndoA2 was co-expressed with EGFP-tagged wild-type NHSL1-F1, specified EGFP-tagged NHSL1-F1 mutants or EGFP alone in HEK293 cells. Proteins were pulled down from lysates using a nanobody against EGFP and blots were probed using antibodies against Myc and EGFP. Blot representative of three independent experiments. (**E**) Quantification of band intensity from **D**. Bands from α-Myc panel were normalised to the corresponding band from α-EGFP panel. Each value was then normalised to that of wild-type NHSL1-F1 which was set to 100%. Results represent means ± SEM with individual data points from each replicate shown. N_=_3 independent biological experiments. Statistics performed using one-way ANOVA with Dunnett’s multiple comparisons test: **P = 0.001. ****P<0.0001.

As NHSL1 and EndoA1 can both interact with VASP^13^, we speculated that NHSL1 may be able to interact with EndoA2 indirectly via VASP, in addition to its ability to interact with the SH3 domain of EndoA2 directly. To account for this possibility, we generated a mutant NHSL1-F1 construct in which all 11 EndoA2 binding sites as well as both FP4 motifs were mutated (NHSL1*EndoAbiFP4*, Fig. 5A). We next aimed to evaluate the ability of all previously described NHSL1-F1 mutants (NHSL1*Endo*, NHSL1*EndoAbi*, NHSL1*FP4*, NHSL1*EndoAbiFP4* and NHSL1*Abi*) to interact with full length EndoA2 in cells. We co-expressed EGFP-tagged wild-type or mutant NHSL1-F1 constructs with Myc-tagged EndoA2 in HEK293 cells. We found that all five NHSL1 mutants showed significant reduction in binding to EndoA2 compared to wild-type NHSL1 (Fig. 5D, 5E). We saw roughly a 5-fold reduction in binding of EndoA2 to NHSL1*Endo*, NHSL1*EndoAbi*, NHSL1*EndoAbiFP4* and NHSL1*Abi*, while NHSL1*FP4* showed an approximate 2-fold reduction. These findings suggest that EndoA2 binds to multiple SH3-domain binding sites in NHSL1 and can additionally interact indirectly via Ena/VASP proteins.

### Evaluation of FEME activity in the MDA-MB-231 triple negative breast cancer cell line

To accurately quantify FEME carrier formation in cells, we developed a defined analysis pipeline to quantify the number of FEME carriers within the whole lamellipodium but excluding the very edge where FEME priming patches reside. The quantified number of FEME carriers per cell was divided by the total area of the lamellipodium to control for cell size. We first tested this analysis pipeline on a dataset generated in unstimulated BSC1 cells (Fig. S5A), those stimulated with a total of 20% foetal calf serum (FCS) (Fig. S5B) or those stimulated with 10 ng/ml EGF (Fig. S5C). Before quantifying the number of FEME carriers per cell, we first assessed the percentage of cells in each condition where FEME priming patches and carriers were detectable. These values estimated the proportion of cells with active FEME within each experimental group. FEME priming patches could be observed in approximately 65% of unstimulated BSC1 cells (Fig. S5D), with around 31% of cells displaying FEME carriers (Fig. S5E). Notably, stimulation with FCS or EGF did not significantly increase these percentages (Fig. S5D,E). Employing our FEME carrier quantification pipeline, we found no significant difference in carrier formation between cells under resting conditions and cells stimulated with 10% extra serum as reported in^7^, but this may have been due to differences in growth factor content of different FCS batches (Fig. S5F). As expected, we observed a significant increase in the number of carriers present in cells stimulated with EGF from 3.37 ± 0.33 to 5.09 ± 0.44 (mean ± SEM, Fig. S5F), in agreement with previous reports^5^.

EndoA2 depletion impairs the uptake of EGFR (ERBB1) in triple-negative breast cancer cells (absence of oestrogen-, progesterone-, and HER2 receptors), and its family member HER2 (ERBB2) in HER2-positive cells^36,37^. Furthermore, EndoA2 and multiple FEME priming proteins have been shown to promote invasion and metastasis of breast cancer and correlate with poor patient prognosis^27,36–40^. Importantly, whilst active at variable levels in different cell types^7^, FEME has not been studied in the context of breast cancer. Therefore, we aimed to quantify levels of FEME in MDA-MB-231 cells, chosen for the characterised role of EndoA2 in EGFR uptake, invasion, and metastasis in this cell line^36^. We evaluated expression levels by western blot and found that NHSL1 was expressed at a higher level in MDA-MB-231 compared to B16-F1 and BSC1 cells but at a lower level compared to HUVEC cells. In contrast, NHSL2 was not expressed in MDA-MB-231 (Fig. S1B). EndoA2 was comparably expressed in BSC1 and MDA-MB-231 cells (Fig. S1B). We stained unstimulated cells (Fig. S5G), those stimulated with 10% extra FCS (20% total, Fig. S5H) and those stimulated with 10 ng/ml EGF (Fig. S5I) for EndoA2. We found that 74% of unstimulated MDA-MB-231 cells contained FEME priming patches (Fig. S5J), with 45% of cells containing FEME carriers (Fig. S5K). As with BSC1 cells, the percentage of cells containing FEME priming patches or FEME carriers was not significantly increased by FCS or EGF stimulation (Fig. S5J,K). Whilst FCS stimulation also had no effect on the number of carriers per cell, stimulation with EGF resulted in a change in the number of FEME carriers in MDA-MB-231 cells from 4.19 ± 0.65 to 5.49 ± 0.64 (mean ± SEM) which was not statistically significant (Fig. S5L). Prior starvation did not potentiate the effect of stimulation on FEME carrier formation in either cell line (Fig. S6). Importantly, MDA-MB-231 cells displayed numbers of FEME carriers comparable to that of the BSC1 cell line, a documented cellular model to study FEME, across all three conditions (Fig. S5F,L).

### NHSL1 promotes FEME through interactions with EndoA2 and Ena/VASP proteins

To explore the function of NHSL1 in FEME we generated NHSL1 knock-out and knock-down MDA-MB-231 cell lines through CRISPR-Cas9 technology and stable expression of shRNAs targeting all isoforms of NHSL1 respectively. Using RT-qPCR, we confirmed successful knock-down of all NHSL1 isoforms in two stable MDA-MB-231 cell lines each expressing a separate shRNA against all isoforms of NHSL1, relative to a control cell expressing a non-targeting shRNA (Fig. S7A). For NHSL1 knock-out cells, we extracted the whole genome from potential clones and sequenced around the intended cut site in exon 2 common to all NHSL1 isoforms. We used the webtool DECODR (https://decodr.org/) to confirm genomic knock-out of NHSL1 in candidate clones^41^. DECODR can be used to align an experimental sequencing file (knock-out clone) to a control sequencing file (wild-type cell) and identify indels present amongst alleles in the experimental sequence. We identified two individual clones (NHSL1 KO 1 and NHSL1 KO 2) in which all alleles contained frame shift mutations resulting in premature termination codons (Fig. S7F,G). We verified loss of NHSL1 expression in the NHSL1 KO clones by western blot (Fig. S7B). Knockout of NHSL1 did not result in compensatory changes in NHSL2 expression nor in changes to EndoA2 expression (Fig. S7C,D).

To assess the effect of NHSL1 depletion on FEME, we stained our control (Fig. 6A) and NHSL1 knock-down cells (Fig. 6B, C) for EndoA2. Cells were stimulated with 10 ng/ml EGF in all cases to maximise the number of FEME carriers observed. EndoA2 staining revealed that the proportion of cells containing FEME priming patches or FEME carriers was not significantly different between knock-down and control cells (Fig. S8A,B). Importantly, we found that in both NHSL1 knock-down cell lines, the number of FEME carriers was significantly reduced compared to the control line (Fig. 6D).

**Figure 6.**
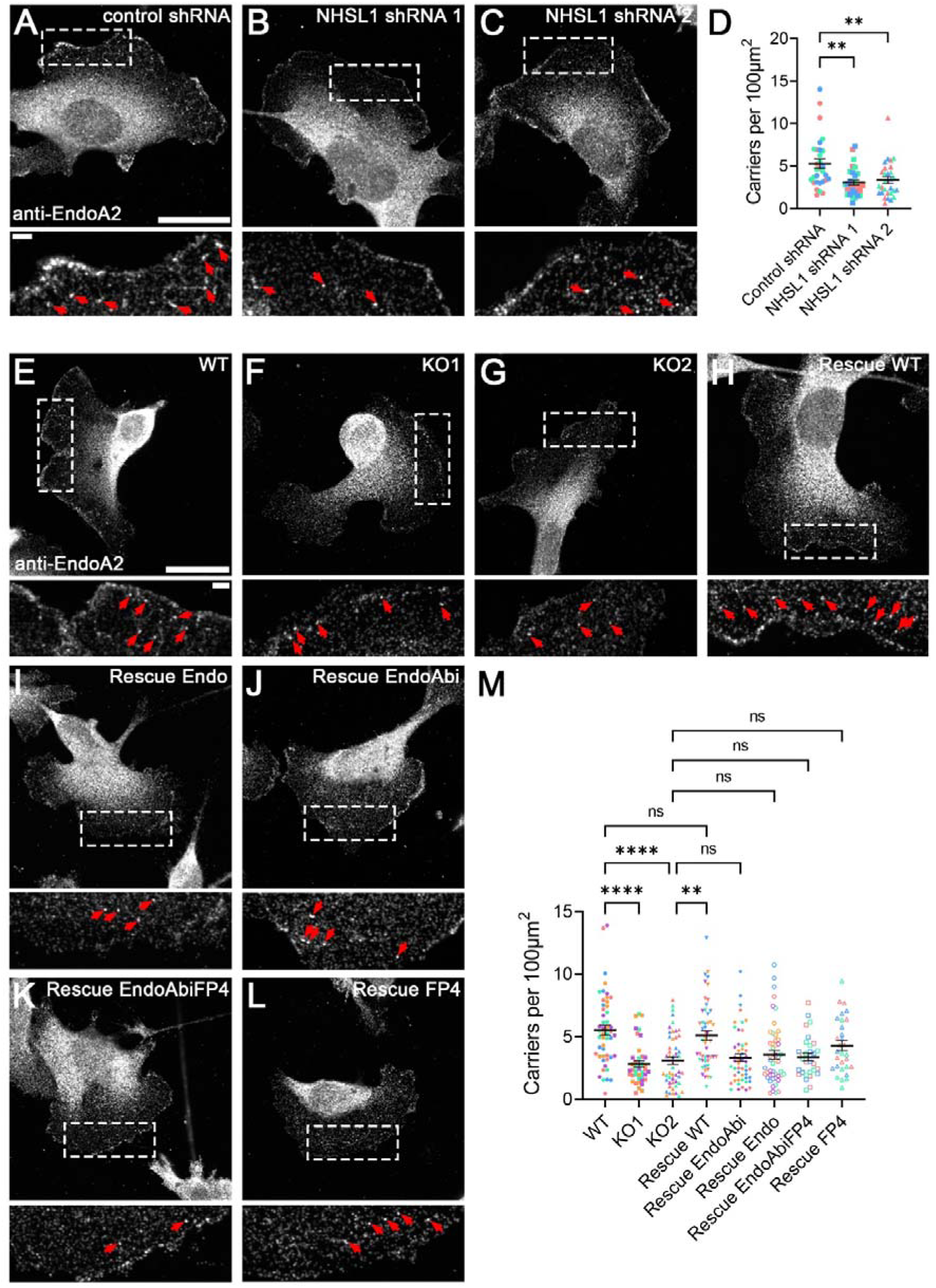
NHSL1 promotes FEME through interactions with EndoA2 and the Ena/VASP proteins. (**A** – **C**) MDA-MB-231 cells stably expressing a control shRNA (**A**) or shRNAs against all NHSL1 isoforms (**B**, **C**) were grown in normal growth medium on collagen and then stimulated with media containing 10 ng/ml EGF. Cells were fixed, immuno-stained for EndoA2 and imaged at a single confocal plane placed in the middle (in Z) of the lamellipodium. Scale bar: 10 μm, Scale bar of inset: 2 μm. Red arrows mark FEME carriers. (**D**) Quantification of the number of FEME carriers per cell normalised to a 100 μm^2^ area. Data represents means ± SEM with individual data points shown. Each data point represents 1 cell. Colours represent individual biological replicates. N = 30 cells per condition across 3 biological replicates. Statistics performed using Kruskal-Wallis test, ** p<0.008. (**E** – **L**) Wild-type (WT), (E) NHSL1 KO 1 (**F**) and KO 2 (**G**) MDA-MB-231 cells were transfected with Myc-only control plasmid. NHSL1 KO 2 cells were also transfected with wild-type NHSL1-F1 (**H**) and indicated mutant NHSL1-F1 constructs (**I**-**L**). Following blasticidin selection, cells were grown in normal growth medium on collagen and then stimulated with media containing 10ng/ml EGF. Cells were fixed, immuno-stained for EndoA2 and imaged at a single confocal plane placed in the middle (in Z) of the lamellipodium. Scale bar: 10 μm, Scale bar of inset: 2 μm. Red arrows mark FEME carriers. (**M**) Quantification of the number of FEME carriers per cell normalised to a 100 μm^2^ area. Data represents means ± SEM with individual data points shown. Each data point represents 1 cell. Colours represent individual biological replicates. N = 30-50 cells per condition across 3-5 biological replicates. Statistics performed using Kruskal-Wallis test, **** p<0.0001, ** p=0.0014, ns p>0.05.

To independently confirm the knock-down result and to investigate whether the interaction of NHSL1 with EndoA2 or Ena/VASP proteins is required for its role in FEME, we employed a knock-out-and-rescue approach, using our two NHSL1 knock-out MDA-MB-231 lines (NHSL1 KO 1 and NHSL1 KO 2) and wild-type MDA-MB-231 cells. Expression of NHSL1 was rescued in the NHSL1 KO 2 line with Myc-tagged wild-type NHSL1-F1 or one of four Myc-tagged mutant constructs (NHSL1*Endo*, NHSL1*EndoAbi*, NHSL1*EndoAbiFP4*, NHSL1*FP4*, Fig 5A). All constructs conferred blasticidin resistance, and cells were selected with blasticidin 24 hours post-transfection to ensure that all analysed cells re-expressed Myc-NHSL1-F1. Following selection, cells were stimulated with EGF and stained for EndoA2 to assess the presence of FEME carriers and priming patches (Fig. 6E-L). Knocking out NHSL1 reduced the percentage of cells with FEME priming patches but this reduction did not reach statistical significance (Fig. S8C). However, NHSL1 knock-out caused a significant reduction in the percentage of cells containing FEME carriers (Fig. S8D). Re-expression of wild-type NHSL1, but not mutant constructs, restored the percentage of cells which exhibited FEME priming patches and carriers to near wild-type levels, albeit this change was not statistically significant (Fig. S8C, D).

Consistent with our data in NHSL1 knock-down cells, we observed a significant reduction in the number of FEME carriers present in both NHSL1 knock-out cell lines compared to wild-type (Fig. 6M). Furthermore, this phenotype was completely rescued in NHSL1 KO 2 cells through re-expression of wild-type NHSL1 (Fig. 6M). However, none of the EndoA2 or Ena/VASP binding mutant NHSL1-F1 constructs were able to significantly restore the number of FEME carriers back to wild-type levels in NHSL1 KO 2 cells (Fig. 6M). Taken together, these data suggest that NHSL1 supports the invagination and scission of FEME priming patches into FEME carriers and that the interaction of NHSL1 with EndoA2 mediates its function in regulating FEME carrier biogenesis. Moreover, the data suggest that the interaction of NHSL1 with Ena/VASP proteins is also important for this process, potentially through regulation of actin dynamics at sites of endocytosis.

### NHSL1 is not essential for dynamin recruitment but promotes F-actin polymerisation at FEME sites

Finally, we asked whether NHSL1 may support the invagination and scission of FEME priming patches into FEME carriers by increasing F-actin polymerisation or by recruiting dynamin. Immunofluorescence analysis and quantification of dynamin localisation or F-actin intensity at sites of FEME revealed that NHSL1 knockout did not affect dynamin recruitment (Fig. 7 A-D) but significantly impaired F-actin polymerisation (Fig. 7 E-H). Taken together this suggests that NHSL1 does not recruit dynamin but instead functions to increase actin polymerisation to support FEME carrier formation.

**Figure 7.**
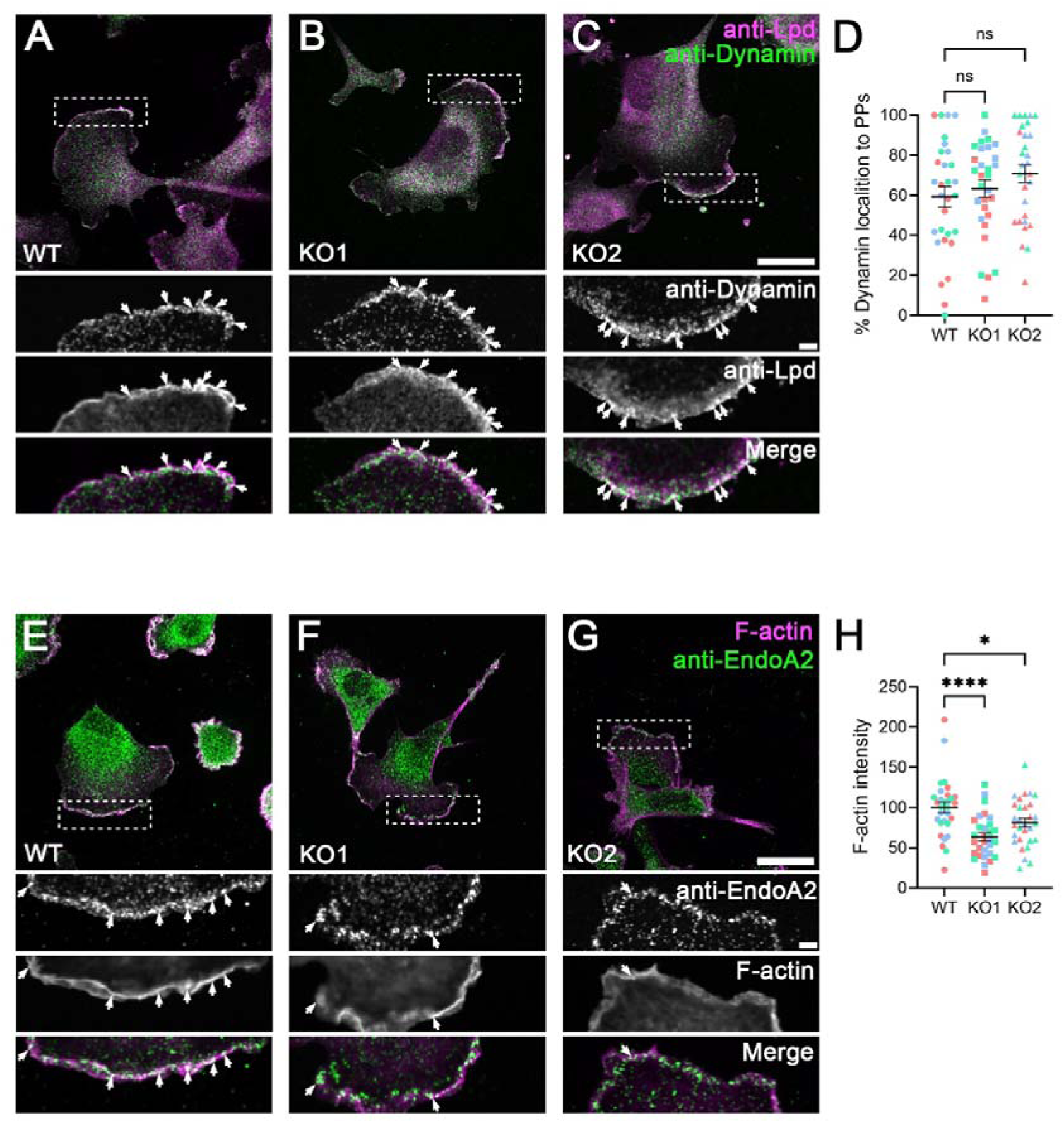
NHSL1 promotes actin polymerisation at FEME sites and is dispensable for dynamin recruitment. (**A** – **C**) Wild-type (WT) (**A**), NHSL1 KO 1 (**B**) and KO 2 (**C**) MDA-MB-231 cells were grown in normal growth medium on collagen. Cells were fixed, stained for Lamellipodin (Lpd) and Dynamin1/2, and imaged at a single plane placed in the middle (in Z) of the lamellipodium. Scale bar: 20 μm, Scale bar of inset: 2 μm. White arrows mark FEME priming patches to which Dynamin1/2 localises. (**D**) Quantification of the percentage of FEME priming patches to which Dynamin1/2 localises. Data represents means ± SEM with individual data points shown. Each data point represents 1 cell. Colours represent individual biological replicates. N = 30 cells per condition across 3 biological replicates. Statistics performed using one way ANOVA with Dunnett’s multiple comparisons test: ns p>0.05. (**E** – **G**) Wild-type (WT) (**E**), NHSL1 KO 1 (**F**) and KO 2 (**G**) MDA-MB-231 cells were grown in normal growth medium on collagen. Cells were fixed, stained for EndoA2 and F-actin (alexa647-phalloidin), and imaged at a single plane placed in the middle (in Z) of the lamellipodium. Scale bar: 20 μm, Scale bar of inset: 2 μm. White arrows mark FEME priming patches to which F-actin localises. (**H**) Quantification of F-actin intensity at FEME priming patches, normalised to the mean fluorescence intensity in WT cells which is set to 100. Data represents means ± SEM with individual data points shown. Each data point represents 1 cell. Colours represent individual biological replicates. N = 30 cells per condition across 3 biological replicates. Statistics performed using one way ANOVA with Dunnett’s multiple comparisons test: **** p<0.0001, ** p<0.05.

## Discussion

Despite insights into the molecular mechanism driving FEME priming patch formation, the process underpinning FEME carrier formation is poorly defined. Whilst actin polymerisation is known to be essential for FEME^5^, how it is controlled to support FEME carrier formation is unclear. Here we report that both NHSL1 and its family member NHSL2 co-localise and engage in direct, multivalent interactions with EndoA2, a key regulator of FEME. In addition, we show that NHSL1 co-localises with and binds directly to all three Ena/VASP proteins via their EVH1 domain. We reveal that NHSL1 supports FEME carrier formation and that this function is dependent on its interaction with EndoA and Ena/VASP proteins. Moreover, we found that NHSL1 is not required for dynamin recruitment but instead increases F-actin polymerisation at sites of FEME. This suggests that NHSL1 promotes FEME through EndoA2 mediated membrane bending and Ena/VASP mediated actin polymerisation.

In BSC1 or HUVEC cells, established cellular models of FEME, we observed NHSL1 or NHSL2 at approximately half of FEME priming patches. As NHSL1/2 do not localise to all FEME priming patches, this suggests that NHSL1/2 assemble at FEME priming patches after EndoA2 is initially recruited. Thus, NHSL1/2 may support further EndoA2 recruitment and clustering at FEME priming patches after their formation and throughout their maturation into FEME carriers. Importantly, by staining for EndoA2 we cannot distinguish nascent FEME priming patches from those which contain activated receptors and are starting to invaginate. Therefore, the presence or absence of NHSL1/2 may reflect the level of maturity of a given FEME priming patch. The exact mechanism by which NHSL1/2 itself is recruited to FEME priming patches remains unclear but may require its interaction with the small GTPase Rac^29^.

As NHSL1/2 were also found at a small proportion of FEME carriers, it is likely that they remain associated during vesicle scission, dissociating from the FEME carrier shortly thereafter. However, in B16-F1 cells, NHSL1 and NHSL2 mostly co-localised with EndoA2 at internalised carriers, with minimal co-localisation at the plasma membrane, likely due to the absence of EndoA2-positive priming patches observed in this cell line. This lack of FEME priming patches could be explained by high levels of constitutive FEME in B16-F1 cells, driven by high Rac activity, as Rac is essential for FEME^5^ and B16-F1 cells are Rac dominated, as evidenced by their propensity to form large Rac-driven lamellipodia.

Like Lpd^12^, both NHSL1 and NHSL2 harbour multiple binding sites for the SH3 domain of EndoA2. Interestingly, only three of these sites are conserved in both proteins: the two previously characterised Abi binding sites (Sites 6 & 9 in NHSL2 – Fig. 3F, Site 4 & 6 in NHSL1 – Fig. 3C) and one further site (Site 13 in NHSL2 – Fig. 3F, Site 9 in NHSL1 – Fig. 3C). Despite low amino acid sequence conservation among NHS proteins^29^, this is consistent with a high conservation of biochemical function. Mutating the EndoA2 binding sites significantly reduced but did not completely block binding of EndoA2 to NHSL1 in pull-down experiments. We propose two potential explanations for this finding. Firstly, it is possible that other proteins may facilitate an indirect interaction of NHSL1 with EndoA2. Mutating the Ena/VASP binding sites was able to partially disrupt binding of EndoA2 to NHSL1, indicating that the interaction between NHSL1 and EndoA2 is partially indirect. However, introducing these mutations into EndoA2 binding mutants of NHSL1 did not further impair the interaction of these mutants with EndoA2, suggesting that further indirect binding partners of both proteins exist. Secondly, EndoA2 may bind to other non-canonical SH3-domain binding sites in NHSL1 that were not identified in this study. In support of this, Teyra and colleagues found that the SH3 domain of EndoA2 can bind to atypical binding motifs which lack proline residues altogether^17^. Interestingly, we observed interaction of NHSL1/2 with the SH3 domains of EndoA1 and EndoA3 in addition to EndoA2. As EndoA1 and EndoA3 are implicated in other clathrin-independent endocytosis (CIE) pathways^9,42–44^, this opens the possibility that NHSL1/2 also support these CIE mechanisms.

We observed that the shared EndoA2 and Abi binding sites in NHSL1 mediated the strongest interaction with EndoA2 *in vitro* (Fig. 3) but in cells the sites solely mediating interaction with EndoA2 (Fig. 5) were equally important. Consistent with previous studies, this finding underscores the binding promiscuity of the EndoA SH3 domain which may be crucial for multivalent interactions of EndoA with its various binding partners. Multivalent interactions of EndoA1 with the C-terminus of Lpd and the third intracellular loop of the β1-AR have been shown to be sufficient to trigger biomolecular condensate formation or liquid-liquid phase separation (LLPS) of EndoA with these proteins^45^. Therefore, the multivalent interactions of EndoA2 with NHSL1 and NHSL2 may similarly drive the formation of EndoA2- and NHSL1/2-containing condensates through LLPS which may contribute to FEME priming patch formation and/or transformation of condensate associated membranes into endocytic buds. However, how condensates containing EndoA2, Lpd, and NHSL1 behave needs to be explored.

The key findings of this study are that NHSL1 depletion impairs FEME and F-actin polymerisation at FEME sites, suggesting that NHSL1 supports F-actin polymerisation driving the invagination of FEME priming patches and scission into FEME carriers. Importantly, our data suggests that the interaction of NHSL1 with EndoA2 drives FEME carrier biogenesis and this may be facilitated by LLPS. NHSL1 may take over from Lpd after receptor activation to recruit and cluster EndoA2, thereby activating it to support invagination and/or scission. We also found that NHSL2 interacts and co-localises with EndoA2, suggesting that it may play a similar role to NHSL1 in FEME. However, NHSL2 expression is more tissue-restricted than that of NHSL1, suggesting that its involvement in FEME may be cell type dependent. In line with the strong impact of NHSL1 depletion on FEME in the MDA-MB-231 cell line, NHSL2 mRNA levels are reported to be very low in this cell type, according to the Human Protein Atlas ^46^ and in western blots its expression was not detectable (Fig. S1B).

We found that NHSL1 co-localises with VASP and EVL at vesicles which may represent FEME carriers and with all three Ena/VASP proteins at the very edge of lamellipodia. We showed that NHSL1 interacts directly with all three Ena/VASP proteins via their EVH1 domain. Our data supports previous findings by Hwang et al, which identified an NHSL1 peptide containing both FP4 motifs as a high affinity Mena EVH1 ligand in a bacterial peptide-display screen^47^. We showed that the two NHSL1 FP4 motifs individually interact with VASP (Fig. 4E). Similarly, Hwang et al. found that each motif can bind the Mena EVH1 domain individually^47^. However, they also reported that a higher binding affinity is achieved through simultaneous binding of both FP4 motifs to a single EVH1 domain^47^. This enhanced affinity is comparable to that of the *L. monocytogenes* ActA protein for the Mena EVH1 domain, which represents one of the strongest characterised EVH1 ligands^47^.

Biomolecular condensates formed on lipid bilayers from Lpd and EndoA1 have been shown to recruit and cluster VASP, thereby promoting actin polymerisation within these condensates^13^. Likewise, NHSL1 and EndoA2 may form condensates at the plasma membrane to drive actin polymerisation at FEME priming patches through Ena/VASP protein recruitment. In support of this, we found that the interaction of NHSL1 with Ena/VASP proteins is required for its function in FEME. However, exactly how NHSL1-EndoA-VASP condensates may differ from Lpd-EndoA-VASP condensates and how the transition from an Lpd-EndoA condensate to a NHSL1-EndoA condensate may occur needs to be explored.

We previously found that endogenous Mena and VASP localise strongly to FEME priming patches^13^, suggesting a function in FEME membrane invagination and/or carrier scission. Lpd, required for EndoA recruitment to FEME priming patches, also binds and recruits Ena/VASP proteins to subcellular locations such as lamellipodia and clathrin-coated pits^12,26^. Therefore, Lpd and NHSL1 may both recruit Ena/VASP proteins to sites of FEME. Intriguingly, the interaction of Lpd with VASP and Mena is negatively regulated by GSK3^48^, a known inhibitor of FEME^7^. This may suggest that inhibition of local GSK3 activity may be required to initiate FEME carrier formation by promoting the interaction of Lpd with VASP and Mena.

Whilst actin nucleation at sites of FEME is mediated by the Arp2/3 complex^5^, the mechanisms controlling actin filament elongation in FEME were previously unclear. Here, we show that NHSL1 promotes F-actin polymerisation at FEME sites and that interaction with Ena/VASP proteins supports FEME. Our findings thus indicate that Ena/VASP-mediated actin filament elongation may provide the force for plasma membrane invagination and or scission. Similarly, we previously showed that Lpd recruits the Ena/VASP protein, Mena, to mediate the scission step of clathrin-mediated endocytosis of the EGF receptor^12^.

In summary, this study reveals a key role for NHSL1 in FEME which depends on its interactions with membrane bending EndoA and actin elongating Ena/VASP proteins.

## Methods

### Molecular biology, plasmids and reagents

Myc-tagged murine NHSL2 (NM_001163610) was purchased from Origene (MR223518) and was used to clone full length murine NHSL2, and the six NHSL2 fragments (Fig. S2E) into pENTR3C (Invitrogen). Full length human NHSL1-F1 (including exon 1f and excluding exons 3 and 6) was generated by cloning overlapping EST clones Flj35425 and DKFZp686P1949 into pENTR3C^29^. Full length human NHSL1-A1 including Exon 1a and excluding exons 3 and 6 was generated by NEB HIFI assembly using a synthetic DNA fragment (gBlocks^TM^, IDT) containing Exon 1a and overlapping sequences and human NHSL1-F1 cDNA into pENTR3C. NHSL1*FP4(1)* and NHSL1*FP4(1+2)* in pENTR3C were generated by mutating Phenylalanine and Proline residues to Alanine (FP4-1: FLPPPPP>ALAAPPP; FP4-2: VFPPPPP>AAAAPPP) using primer-mediated site-directed mutagenesis followed by DpnI treatment to digest parental, unmodified plasmid. cDNA for NHSL1*Abi* was generated by mutating select Arginine and Proline residues to Alanine by site-directed mutagenesis (see Fig. 5A)^29^. cDNAs for NHSL1*Endo*, NHSL1*EndoAbi*, and NHSL1*EndoAbiFP4* were generated using HiFi assembly (New England Biolabs) of eight overlapping DNA fragments, each containing mutations to EndoA2 binding sites at overlaps, into pENTR3C. To mutate each EndoA2 binding site, every Proline residue within that site was substituted for an Alanine (see Fig. 5A). For NHSL1*Endo*, six of these fragments were amplified by PCR and for the other two fragments pre-synthesised short double stranded cDNAs (gBlocks^TM^, IDT) were used. For NHSL1*EndoAbi* and NHSL1*EndoAbiFP4*, five of the fragments were amplified by PCR and the other three fragments were pre-synthesised short double stranded cDNAs (gBlocks^TM^, IDT). All PCR-amplified fragments used wild-type full length NHSL1-F1 as a template, with the exception of the fragment in NHSL1*EndoAbiFP4* containing the two mutated FP4 motifs which instead used NHSL1*FP4(1+2)*. Constructs for N- or C-terminally EGFP-, mScarlet- or Myc-tagged NHSL1 or NHSL2 were generated by transferring cDNAs in pENTR3C into the following DEST vectors using Gateway^®^ recombination (Invitrogen): pCAG-EGFP-DEST, pCAG-DEST-mScarlet-I, pCAG-Myc-DEST-IRES-Blast. Constructs for N-terminally GST-tagged NHSL2 fragments were generated by transferring cDNAs in pENTR3C into pDEST15 (Invitrogen) using Gateway^®^ recombination (Invitrogen). Constructs for N-terminally GST-tagged NHSL1 fragments (Fig. 3A, S2B) were generated by amplifying fragments from full length NHSL1-F1 and cloning these into pGEX-6P1 (Cytiva).

Human full length endophilin-A1 (EndoA1, SH3GL2) in pDONR201 (HSCD00000899, Harvard Institute of Proteomics) and human endophilin-A3 (EndoA2, SH3GL3, IMAGE5197246-AK68-m23) in pENTR11 (Invitrogen) were transferred into pCMV-DEST-EGFP using Gateway^®^ recombination (Invitrogen)^12^. C-terminally EGFP-tagged rat full length endophilin-A2 (EndoA2, SH3GL1) was a kind gift from Dr Pietro De Camilli (Yale University, USA). The SH3 domains of EndoA1-3 were amplified by PCR from full length constructs and cloned into pGEX-6P1 (Cytiva, GST-tagging), and the EndoA2 SH3 domain was cloned into pMAL-c2g (New England Biolabs, MBP-tagging)^12^. Full length EndoA2 and EndoA2Δ*SH3* (aa 1-311) were cloned into pENTR3C and transferred into pRK5-Myc-DEST using Gateway recombination (Invitrogen).

Murine VASP, Mena, and EVL cDNAs (kindly gifted by Frank Gertler, MIT) were cloned into pENTR3C (Invitrogen) and transferred into Myc-tagged and mScarlet-tagged mammalian expression vectors using Gateway recombination. EVH1 domains of VASP, Mena and EVL and PRR and EVH2 domains of EVL were amplified from full length constructs, cloned into pENTR3C and transferred into pDEST15 (Invitrogen) using Gateway recombination (Invitrogen).

### Antibodies

#### Primary antibodies

Rabbit polyclonal anti-NHSL1 serum clone #4457 (Eurogentec, ^29^), IF (1:400). Rabbit polyclonal anti-NHSL1 (Atlas Antibodies, HPA029967), WB (1:1000). Rabbit polyclonal anti-NHSL2 (Atlas Antibodies, HPA075115) WB (1:1000), IF (1:100). Mouse monoclonal anti-EndoA2 clone A-11 (Santa Cruz Biotechnology, sc-365704), WB (1:2000), IF (1:200). Rabbit polyclonal anti-Lpd serum clone #3917^26^. Mouse monoclonal anti-Dynamin clone 41 recognising both Dynamin 1 and 2 (BD Biosciences, 610245), IF (1:100). Rabbit polyclonal anti-EVL serum clone #1404, ^49^), WB (1:5000). Mouse monoclonal anti-EGFP (Roche, 11814460001), WB (1:2000). Mouse monoclonal anti-Myc (Sigma Aldrich, M5546), WB (1:5000). Mouse monoclonal anti-MBP (New England Biolabs, E8032S), WB (1:10000). Mouse monoclonal anti-α-tubulin (Sigma Aldrich, T6199), WB (1:5000). HRP-conjugated rabbit monoclonal anti-GAPDH clone 14C10 (Cell Signalling Technology, #3684), WB (1:1000).

#### Secondary antibodies

HRP-goat anti-mouse IgG (Agilent, P0447), WB (1:2000). HRP-donkey anti-mouse IgG (Jackson ImmunoResearch, 715-035-150), WB (1:10000). HRP-goat anti-rabbit IgG (Agilent, P0448), WB (1:2000). HRP-donkey anti-rabbit IgG (Jackson ImmunoResearch, 711-035-152), WB (1:10000). Alexa488-goat anti-mouse IgG (Invitrogen, A11011), IF (1:500). Alexa488plus-goat anti-mouse IgG (Invitrogen, A32723), IF (1:500). Alexa568-goat anti-mouse IgG (Invitrogen, A11001), IF (1:500). Alexa568-goat anti-mouse IgG (Abcam, ab175471), IF (1:500).

#### Other Staining Reagents

Alexa647-Phalloidin (Invitrogen, A30107), IF 1:500.

### Cell culture and transfection

HEK293FT (referred to in this paper as HEK293, ThermoFisher Scientific R70007), B16-F1 (ATCC CRL-6323), BSC1 (Gift from Emmanuel Boucrot, UCL; ATCC CCL-26) & MDA-MB-231 (Gift from Anne Ridley, KCL University of Bristol; ATCC CRM-HTB-26) cells were cultured in high glucose Dulbecco’s Modified Eagle’s Medium (DMEM) containing 2mM L-Glutamine, 1% Penicillin/Streptomycin and 10% foetal bovine serum at 37°C and 10% CO_2_.

Human Vascular Endothelial Cells (HUVEC, Promocell) were cultured in EGM-2 BulletKit™ Medium (Lonza, CC-3162) at 37°C and 5% CO_2_.

HEK293, BSC1, MDA-MB-231 cells and HUVECs were passaged once 90% confluent (approximately every three days) at a ratio of 1:5. B16F1 cells were passaged every two to three days at a ratio of 1:20-1:50.

HEK293 cells were either transfected using 2mg/ml branched polyethyleneimine (PEI, Sigma Aldrich) or Lipofectamine 2000 (Invitrogen). When using PEI, 4µg plasmid DNA and 8µl PEI were diluted separately in 100µl OptiMEM (Gibco), before being mixed together and then incubated at room temperature for 20 min. This mixture was then added dropwise to cells. Cells were incubated for a further 24 hours in normal growth conditions before lysis for western blot experiments. For the pulldown experiments showing an interaction of NHSL1 with VASP, Mena and EVL (Fig. 4), 5μl of Lipofectamine 2000 was diluted in 125 µl of OptiMEM and incubated for 5 min. This was then added to 2 µg of DNA diluted in 125 ul OptiMEM. The DNA and Lipofectamine mix was incubated for 20 min at room temperature. This mixture was then added dropwise to cells.

B16-F1 cells were transfected with X-tremeGENE™ 9 or X-tremeGENE™ 360 (Roche) following manufacturer’s instructions and media was changed 4-6 hours after transfection.

MDA-MB-231 cells were transfected using jetOPTIMUS^®^ (Polyplus) following manufacturer’s instructions and media was changed 4-6 hours after transfection.

BSC1 cells were transfected using Lipofectamine 3000 (Invitrogen) following manufacturer’s instructions and media was changed 4-6 hours after transfection.

### Purification of GST- and MBP-tagged proteins

Chemically competent BL21-CodonPlus(DE3)-RP (Agilent) E. coli cells were transformed with the appropriate plasmids for expression of GST- or MBP-tagged proteins. Protein expression was induced with 0.1mM IPTG (Sigma Aldrich) and bacterial cells were lysed using a sonicator at 15W for 3 min. Cellular debris was pelleted by centrifugation for 20 min (18000 × *g* at 4°C) and GST and MBP fusion proteins were purified using glutathione-Sepharose (GE Healthcare) or amylose (New England Biolabs, Inc.) beads. Purified MBP-fusion proteins were eluted using maltose and dialised/concentrated using Vivaspin filters (Sartorius). To confirm successful protein purification, 10µl of purified proteins on beads were boiled in sample buffer (1.75g DTT, 2.5g SDS, 10ml 1M Tris-base (pH6.8), 12.5ml Glycerol, 750 ul Bromophenol blue in dH2O (50ml total volume)) for 5 min at 95°C and separated by SDS-PAGE. Gels stained using Rapid Blue Colloidal Coomassie Stain (Severn Biotech) following manufacturer’s instructions. Concentrations of proteins to be used in the same experiment were standardized by dilution with sample buffer to ensure equivalency.

### Pulldown, immunoprecipitation, and Western blot

For pulldown and immunoprecipitation experiments, cells were lysed in glutathione S-transferase (GST) buffer (50□mM Tris-HCL, pH 7.4, 200□mM NaCl, 1% NP-40, 2□mM MgCl_2_, 10% glycerol, 10mM NaF, 1mM Na3VO4 and EDTA-free protease inhibitor cocktail tablets (Roche)). Lysates were incubated on ice for 15□min and centrifuged for 10 min (17,000 × *g* at 4□°C). Protein concentration was measured using Precision Red Advanced Protein Assay (Cytoskeleton, Inc.) or Pierce™ BCA Protein Assay Kit (ThermoFisher Scientific) following manufacturer’s instructions. For pulldowns, lysates were incubated with purified GST fusion proteins on glutathione-Sepharose beads (GE Healthcare), GFP-trap beads (Chromotek), Myc-trap beads (Chromotek) or GFP-selector beads (NanoTag). Beads were blocked with 1% BSA before incubating with lysates for 1–2□hours at 4°C. For immunoprecipitations, Protein A bead precleared lysates were incubated with primary antibody or non-immune control IgG followed by 1% BSA blocked protein A beads (Pierce) or protein A/G beads (Alpha Diagnostics). Lysates or samples on beads were boiled in sample buffer for 5 min at 95°C and separated by SDS-PAGE. Proteins were transferred onto polyvinylidene fluoride (PVDF; EMD Millipore). Transfers were performed either at 100 V, 350 mA, 50 W for 90 min. Membranes were blocked at 4°C overnight in in TBST (20□mM Tris-base, 150□mM NaCl, 0.1% (v/v) Tween-20, pH7.6) containing 5% milk (w/v). Blots were probed with primary antibodies diluted in TBST + 5% milk for 1 hour at room temperature or overnight at 4°C. Membranes were then washed in TBST, TBST + 0.5M NaCl and TBST + 0.5% Triton-X-100 for 10 min each. Membranes were probed with secondary horseradish peroxidase (HRP)-conjugated antibodies diluted in in TBST + 5% milk for 1 hour at room temperature. Membranes were washed three further times before being developed using a BioRad ECL Chemiluminescence Kit (Bio-Rad Laboratories) and imaged using a BioRad imager using Image Lab software.

For the pulldown experiments showing an interaction of NHSL1 with VASP, Mena and EVL (Fig. 4), lysates were incubated with GFP-trap beads for 2 hours at 4°C. This was then followed by washing beads 3 times with GST buffer. 5% β-mercaptoethanol was added to each sample and boiled at 70 °C for 10 min before running on 4-12% Bis-Tris gradient gels (Invitrogen) with 1 x MOPS buffer (Invitrogen). Gels were run at 115mA, 200V for 50 min. Proteins were then transferred and detected as above.

For **Fig. S1B** and **Fig S7B-E**, B16-F1, BSC1, MDA-MB-231 cells and HUVECs were lysed in RIPA buffer (150 mM NaCl, 50 mM Tris-HCl pH 7.4, 1% NP40, 0.5% sodium deoxycholate (SDC), 0.1% SDS, phosSTOP tablets (Roche), EDTA-free protease inhibitor cocktail tablets (Roche)). Lysates were incubated on ice for 20□min and centrifuged for 15 min (16,000 × *g* at 4□°C). Protein concentration was measured using Pierce™ BCA Protein Assay Kit (ThermoFisher Scientific) following manufacturer’s instructions. Lysates were boiled in sample buffer for 5 min at 95°C and separated by SDS-PAGE. Proteins were transferred onto nitrocellulose membranes (Amersham). Transfers were performed at 350 mA constant current for 3 h. Membranes were blocked for 30 min at room temperature in in TBST (20□mM Tris-base, 150□mM NaCl, 0.1% (v/v) Tween-20, pH7.4) containing 5% milk (w/v). Blots were probed with primary antibodies diluted in TBST + 3% BSA for 1 hour at room temperature or overnight at 4°C. Membranes were then washed thrice in TBST for 10 min each. Membranes were probed with secondary horseradish peroxidase (HRP)-conjugated antibodies diluted in in TBST for 1 hour at room temperature. Membranes were washed three further times before being developed using EZ-ECL solution (Cytiva) and imaged using a G:BOX Chemi XX9 (Syngene).

### Western Blot Quantification

Western blot quantification was performed using ImageLab software (BioRad). The longest exposure time at which there was no band saturation was selected for each set of bands. For quantification of pulldown experiments, the values from experimental bands were normalised to those from the bands, of the same condition, which corresponded to the protein being directly pulled down (e.g. EGFP-tagged proteins in GFP pull-down experiments). For Fig. 2H the band intensities of the anti-Myc pulldown bands were first normalised to the corresponding anti-Myc input band to control for variability in expression levels between constructs, before normalisation to the anti-GFP pulldown band. Normalised values were then expressed relative to that of the control sample for that experiment which was set to 100% in all replicates.

### Far-Western blot and peptide array

For far-western experiments using GST-tagged NHSL1 and NHSL2 fragments, concentration-adjusted bead fractions were boiled in sample buffer, separated by SDS-PAGE and transferred onto a PVDF membrane as described above. For far-western experiments using full length NHSL1 constructs purified from HEK293 cell lysates, Myc-tagged proteins were purified using Myc-trap beads (Chromotek) as described above. Purified proteins were then boiled in sample buffer, separated by SDS-PAGE and transferred onto a PVDF membrane. Custom PepSPOTS^TM^ peptide arrays were purchased from JPT Peptide Technologies. In all cases, membranes were blocked at 4°C overnight with TBST containing 5% BSA (w/v). Membranes were incubated for one hour with purified MBP-EndoA2 or MBP, diluted in TBST containing 5% BSA, to a final concentration of 0.2–5 µg/ml. Membranes were incubated for one hour with an anti-MBP primary antibody, followed by an HRP-linked secondary antibody. Blots were developed using a BioRad ECL Chemiluminescence Kit (Bio-Rad Laboratories) and imaged using a BioRad imager using Image Lab software.

For VASP overlay assays, custom-made immobilised peptide arrays (Cancer Research UK) with 12 amino acid long peptides covering the FP4 motifs of NHSL1 were overlayed with *in vitro* translated ^35^S-labeled full-length VASP. ^35^S-VASP was made by *in vitro* translation using a T3 driven TNT® Coupled Reticulocyte Lysate system (Promega) from murine VASP cDNA in a pBluescript-KSII-plasmid. Purified GST-fusion proteins covering NHSL1, were blotted and overlaid with *in vitro* translated ^35^S-labeled full-length VASP. Bound VASP was detected using a phosphor-imager Typhoon 9200 (Amersham).

### Immunofluorescence and live-cell imaging

For live-imaging experiments, BSC1 and B16-F1 cells were seeded onto uncoated glass-bottom dishes (Ibidi, 81158) or dishes (Ibidi, 81218-200) coated with 25µg/ml laminin (Sigma Aldrich, L2020) respectively. Cells were left to spread and form lamellipodia for 3-4 hours at 37°C and 10% CO_2_ prior to being imaged.

For immunofluorescence BSC1 cells, HUVECs or MDA-MB-231 cells were seeded on uncoated coverslips (BSC1 cells and HUVECs) or coverslips coated with 10µg/ml collagen (MDA-MB-231 cells). Cells were left to spread for 6-8 hours at 37°C and 10% CO_2_. Cells were either stimulated with 10ng/ml EGF diluted in DMEM, DMEM containing an additional 10% FCS (20% FCS total) or left unstimulated. Stimulated cells were incubated for 4 min at 37°C and 10% CO_2_. For Fig. S5,6 6 hours after initial seeding growth medium was removed and replaced with fresh growth medium with or without 10% FBS, prior to stimulation with EGF as above. Cells were then immediately fixed with 4% paraformaldehyde in PHEM buffer (60□mM PIPES, 25□mM HEPES, 10□mM EGTA, 2□mM MgCl_2_, 0.12□M sucrose) or

Dulbecco’s Phosphate Buffered Saline (For. **Fig 1E**, **Fig. 7** and **Fig. S6**) for 20 min at 37°C and 5% or 10% CO_2_. Cells were reconstituted in cytoskeletal TBS (cTBS, 10x: 200mM Tris-HCl, 1.54M NaCl, 20mM EGTA, 20mM MgCl_2_ x 6H_2_O in dH_2_O, pH 7.5) and stored at 4°C overnight. The next day, fixed samples were washed a further two times with 1x cTBS and then quenched with cTBS containing 50mM NH_4_Cl for 10 min at room temperature. Next, samples were simultaneously blocked and permeabilized in cTBS containing 10% normal goat serum (NGS), 10% BSA (w/v) and 0.05% (w/v) Saponin for 1 hour at room temperature. Samples were washed three times in cTBS and incubated with primary antibodies diluted in cTBS containing 1% BSA (w/v) and 0.05% (w/v) Saponin at 4°C overnight. The next day, samples were washed three times in cTBS and incubated with secondary antibodies diluted in cTBS containing 1% BSA (w/v) and 0.05% (w/v) Saponin for 1 hour at room temperature. Samples were washed three times and then mounted on glass slides with FluorSave mounting medium (Sigma Aldrich) or ProLong™ Gold Antifade Mountant (Invitrogen).

For **Fig 1E**, **Fig. S1C** & **D**, **Fig. 7** and **Fig. S6** cells were imaged using an Olympus SpinSR system with an IX83 microscope (Olympus) two Hamamatsu ORCA-Fusion Cameras and a 100X UPlanApo 1.50 Oil HR lens (Olympus). The system was controlled using Olympus CellSens software. During live-cell imaging, cells were maintained at 37°C and 5% CO_2_ with a cellVivo incubation chamber (PeCon). For fixed imaging, images were taken at a middle confocal plane (i.e. between the top and bottom confocal plain of the lamellipodium) to capture internalised carriers.

For Fig. S3, cells were imaged using an IX 81 microscope (Olympus), with a Solent Scientific incubation chamber, filter wheels (Sutter), an ASI X-Y stage, Cascade II 512B camera (Photometrics), and 60x Plan-Apochromat NA 1.45, or 100× UPlan-Apochromat S NA 1.4 objective lenses controlled by MetaMorph software.

For all other fixed and live-cell imaging experiments, cells were imaged using a CSU-W1 SoRa system (Yokogawa) with an Eclipse Ti-2 inverted microscope (Nikon), two Photometrics Prime 95B sCMOS cameras and a CFI SR HP Apochromat 100X AC oil immersion objective lens (live-cell) or a CFI Plan Apochromat VC 60X water-immersion objective lens with SoRa magnification changer of 2.8x (Fixed). The system was controlled using NIS Elements Software (Nikon). During live-cell imaging, cells were maintained at 37°C and 10% CO_2_ with an incubation chamber (Okolab). For fixed imaging, images were taken at a middle confocal plane (i.e. between the top and bottom confocal plain of the lamellipodium) to capture internalised carriers. All Images were processed using FIJI.

### FEME carrier quantification assay

Wild-type BSC1 cells, wild-type MDA-MB-231 cells and NHSL1 knock-down (and control) MDA-MB-231 cells were seeded, stimulated, fixed and stained for EndoA2 and imaged as described above. For NHSL1 knock-out & rescue experiments, cells were transfected with blasticidin selectable Myc-NHSL1-F or Myc-only control constructs and then selected with 25µg/ml blasticidin for 72 hours. Cells were grown for a further 24 hours in normal growth medium and then were either lysed for western blot analysis or seeded, stimulated, fixed, stained for EndoA2 and imaged as described above. Slides were blinded prior to imaging by covering the sample information with tape and randomly assigning each slide a letter from A onwards depending on the number of conditions.

During each imaging session, every observed non-rounded, spread cell was recorded as either containing at least one FEME priming patch, at least one FEME carrier, at least one of each, or neither. FEME priming patches were defined as EndoA2-positive assemblies (EPAs) at the very edge of lamellipodia and FEME carriers as EPAs at least 1µm from the leading edge, as detected in a confocal plane placed at the middle of the lamellipodium (in Z). For quantification of the number of FEME carriers per cell, images of at least 30 individual cells per condition were taken across at least three separate biological replicates. The number of FEME carriers per cell was quantified in FIJI as follows. Firstly, the quantification area, which encompassed the entire lamellipodium excluding the very edge and the perinuclear area, was drawn by hand. The ‘measure’ tool was then used to measure the area of this region. Next, the ‘3D Objects counter’ plugin was used to simultaneously threshold high intensity EndoA2 spots and count the number of these spots within the selection area. A threshold intensity filter of 40,000 in conjunction with a size filter of 5 pixels was found to most effectively distinguish FEME carriers from background EndoA2 staining. The number of FEME carriers identified by the 3D object counter was divided by the selection area and multiplied by 100 to obtain a normalised FEME carrier count per 100µm^2^ for each cell.

### Quantification of co-localisation at FEME priming patches and FEME carriers

Localisation of proteins of interest to FEME carriers and/or FEME priming patches was assessed using a custom ImageJ macro. Briefly, an initial ROI was drawn around the edge of the lamellipodium to define the area containing FEME priming patches, or around the rest of the lamellipodium to define the area containing FEME carriers. Within these regions, binary masks of FEME priming patches or FEME carriers were generated by intensity thresholding of EndoA2 stain and used to define ROIs. Binary masks of the protein of interest (e.g., NHSL1, NHSL2, Dynamin) were also generated by intensity thresholding and overlap with the priming patch or carrier ROIs was measured. Puncta were classified as positive (i.e., colocalised) if the mean grey value of the mask of the protein of interest within the ROI exceeded 50 out of 255. The percentage of positive ROIs was then calculated for each cell.

### Quantification of F-actin at FEME priming patches

F-actin intensity at FEME priming patches was measured using a custom ImageJ macro. Briefly, an initial ROI was drawn around the edge of the lamellipodium to define the area containing FEME priming patches. Within this region, binary masks of FEME priming patches were generated by intensity thresholding of EndoA2 stain and used to define ROIs. The mean grey value of the phalloidin channel within all priming patch ROIs in each cell was measured to determine F-actin intensity. These values were then normalized to the overall mean intensity in WT cells.

### Generation of NHSL1 knock-down cells

The NHSL1 shRNA constructs (shRNA A: TGCCCATTCTGTGATGATTA shRNA B: TGGATAAATCCCTATCAAGA), Control-shRNA: GCCGATAACCGAGAATACC were cloned into pLL3.7Puro which is derived from pLL3.7 with the addition of a puromycin selection. HEK293 cells were transfected with either of these constructs in combination with pVSVG, pRSV-REV and pMDLg/p plasmids to make shRNA-containing lentiviral vectors. Lentivirus was harvested after 48 hours and used for the transduction of MDA-MB-231 cells, which were subsequently selected with 3µg/ml puromycin for 48 hours.

For RT-qPCR analysis, total RNA was extracted from cells using the RNeasy Mini kit following manufacturer’s instructions (QIAGEN). cDNA was prepared using Superscript IV (Invitrogen) and expression levels of NHSL1 were quantified using standard curves relative to the house keeping gene B2M using gene-specific primers (PrimeTime qPCR primer assays, IDT): hsNHSL1 (Exon 7-8 for the detection of all isoforms):

For: 5’-TAA TGA CCC CCA GTC GAC C-3’, Rev: 5’-AGA ATG GTT TCG GGA GTG G-3’. hsB2M: For: 5’-ACC TCC ATG ATG CTG CTT AC-3’, Rev: 5’-GGA CTG GTC TTT CTA TCT CTT GT-3’.

### Generation of NHSL1 knock-out cells

For knock-out of NHSL1 we used the eSpCas9-plus nuclease. This nuclease is a variant of the *S. pyogenes* Cas9 with increased fidelity, which contains “blackjack” mutations to improve compatibility with 5’-extended sgRNAs^50^. To generate a construct for the simultaneous selection with puromycin, the eSpCas9(1.1) cDNA was removed from the eSpCas9(1.1)-T2A-Puro vector (Addgene, 101039) through restriction digest with AgeI and FseI. At the same time, the cDNA insert of eSpCas9-plus was excised from the eSpCas9-plus plasmid (Addgene, 126767) using the same restriction enzymes. eSpCas9(1.1)-T2A-Puro was a gift from Andrea Németh (Addgene plasmid # 101039 ; http://n2t.net/addgene:101039 ; RRID:Addgene_101039). eSpCas9-plus was a gift from Ervin Welker (Addgene plasmid # 126767 ; http://n2t.net/addgene:126767 ; RRID:Addgene_126767). The vector backbone of eSpCas9(1.1)-T2A-Puro and the eSpCas9-plus insert were purified by gel extraction and ligated together to form the eSpCas9-plus-T2A-Puro plasmid. Three separate webtools (CHOPCHOP - https://chopchop.cbu.uib.no/about, CRISPRon(v1.0) - https://rth.dk/resources/crispr/crispron/ & CRISPick -https://portals.broadinstitute.org/gppx/crispick/public) were used to identify the sgRNA targeting exon 2 of NHSL1, predicted to be the first exon common to all isoforms, with the highest on-target activity. This sgRNA (5’-CAGTCCACTACACTGCCCCG**TGG**-3’) was cloned into eSpCas9-plus-T2A-Puro following a previously described protocol^51^. MDA-MB-231 cells were transfected with the resulting sgRNA-containing plasmid and selected with 3µg/ml puromycin for 48 hours. Clonal cell lines were generated by limited dilution.

To confirm genetic knock-out, genomic DNA was extracted from individual clones using the QIAamp DNA mini kit following manufacturer’s instructions (QIAGEN). Knock-out clones were genotyped by amplifying the genomic region flanking the intended cut site and sequencing the resulting PCR product. Sequencing traces from individual clones were compared to that of the control sample (wild-type MDA-MB-231 cells) using the DECODR webtool (https://decodr.org/)^41^. Two separate clones in which all NHSL1 alleles contained a premature termination codon were selected for use in functional assays. Loss of NHSL1 protein expression was also assessed by western blot.

### Statistical analysis

All statistical analysis was performed using GraphPad Prism software. In all cases, normality of data was first assessed using D’Agostino & Pearson and Shapiro–Wilk normality tests. For figures that contain data that was tested for statistical significance, the statistical and multiple comparisons tests used as well as the relevant *P* values are all stated in the corresponding legend of that figure. *P* values□<□0.05 were considered significant.

## Data availability

All data supporting the findings of this study are available within the paper and its Supplementary Information.

## Acknowledgements

We thank Emmanuel Boucrot (University College London, UK), Pietro De Camilli (Yale University, New Haven, CT, USA), Frank Gertler (MIT, Cambridge, MA, USA), Andrea Németh (University of Oxford, UK), Anne Ridley (Bristol University, UK), and Ervin Welker (Biological Research Centre, Szeged, Hungary) for reagents. Antonio P.A. Ferreira (Harvard Medical School & Brigham and Women’s Hospital, Boston, MA, USA) for advice on the FEME assay. William Barrell (King’s College London, UK) for qPCR training. The King’s College London Nikon Imaging Centre and Babraham Institute Imaging Facility for support in confocal microscopy. J.C. was supported by a Medical Research Council, UK (MRC) studentship. S.J. was supported by a Biological Science Research Council, UK (BBSRC) studentship. This work was supported by grants from the Biotechnology and Biological Science Research Council, UK (BB/N000226/1, BB/R015953/1, M.K.) (BBS/E/B/000C0433, BB/CCG2210/1, H.S.) and an EPSRC Frontier guarantee grant (EP/Z000114/1, H.S.).

