## Supplemental information for "Nance-Horan Syndrome-like 1 interacts with endophilin A2 and Ena/VASP proteins to promote fast endophilin-mediated endocytosis"

**Cope et al. 2025**

**Supplementary Information**

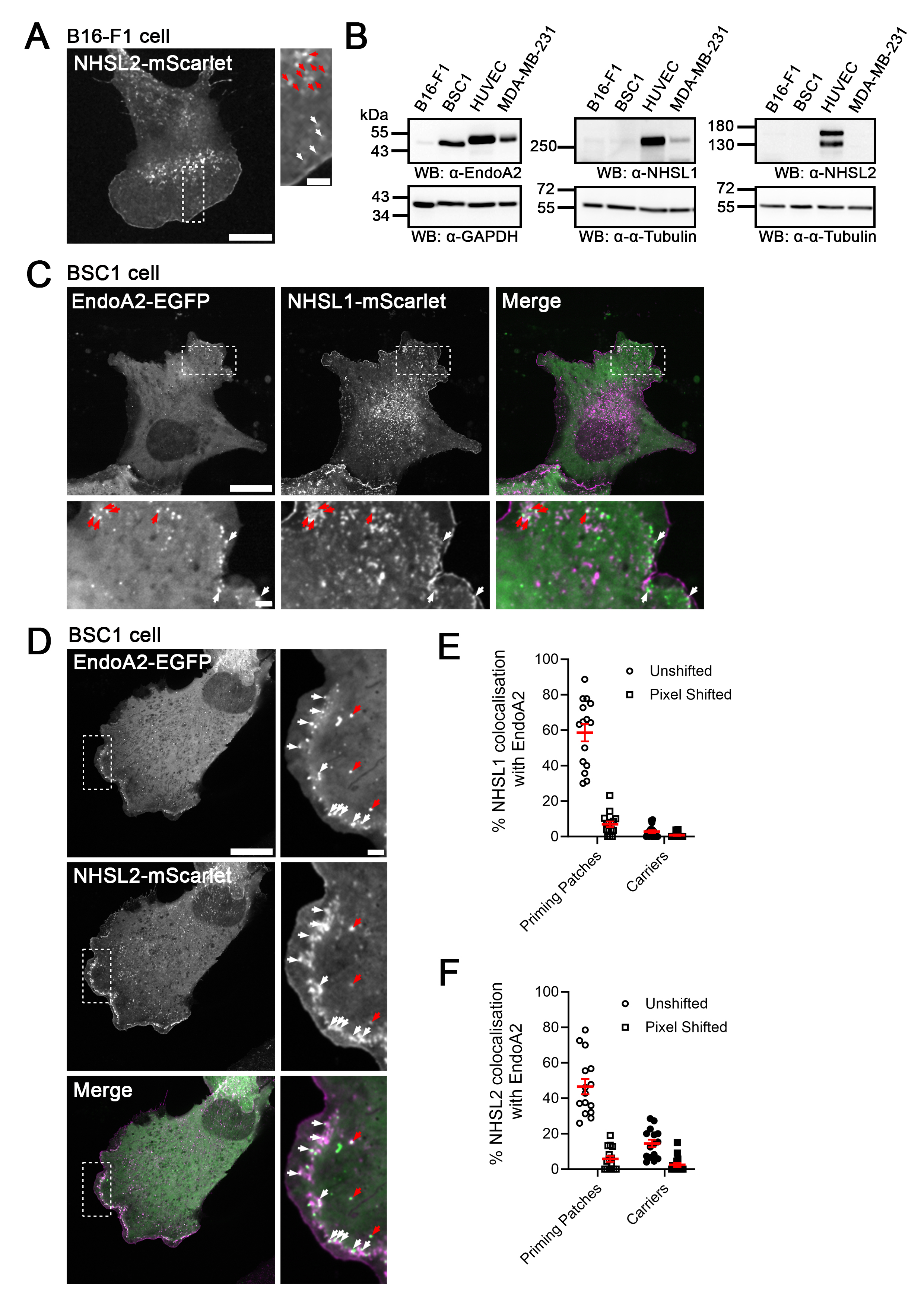

**Figure S1.** *(previous page)* **Related to Figure 1.** **Expression and localisation of NHSL1, NHSL2 and EndoA2 in multiple cell lines.** (**A**) B16-F1 cells were transfected with mScarlet-tagged NHSL2 and plated on laminin. Localisation of tagged proteins was imaged using live confocal microscopy (**Supplementary Movie 1**). White and red arrows mark distinct populations of vesicular puncta. Scale bars: 10µm. Scale bar of insets: 2µm. Images representative of three independent experiments. (**B**) Western blots showing protein levels of EndoA2, NHSL1 and NHSL2 in BSC1, B16-F1, HUVEC and MDA-MB-231 cells. Alpha-tubulin (α-tubulin) and GAPDH were used as loading controls. Blots are representative of three independent experiments. (**C**, **D**) BSC1 cells were co-transfected with EGFP-tagged EndoA2 and mScarlet-tagged NHSL1 (**C**, **Supplementary Movie 4**) or NHSL2 (**D**, **Supplementary Movie 5**). Localisation of tagged proteins was imaged using live confocal microscopy. White and red arrows mark FEME priming patches and carriers respectively. In merged stills, mScarlet-tagged NHSL1 and NHSL2 are represented in magenta, while EGFP-tagged EndoA2 is represented in green. Scale bars: 20µm. Scale bar of insets: 2µm. Images representative of three independent experiments. (**E**, **F**) The percentage of FEME priming patches and carriers at which endogenous NHSL1 (**Fig. 1C**) or NHSL2 (**Fig. 1E**) was quantified before and after shifting the EndoA2 channel by 10 pixels in the X and Y directions. Normalised percentages, calculated by subtracting the pixel shifted values from the unshifted values, are shown in **Fig. 1D, F**. Data represent means ± SEM with individual data points shown. Each data point represents one cell. N = 15 cells across 3 biological replicates.

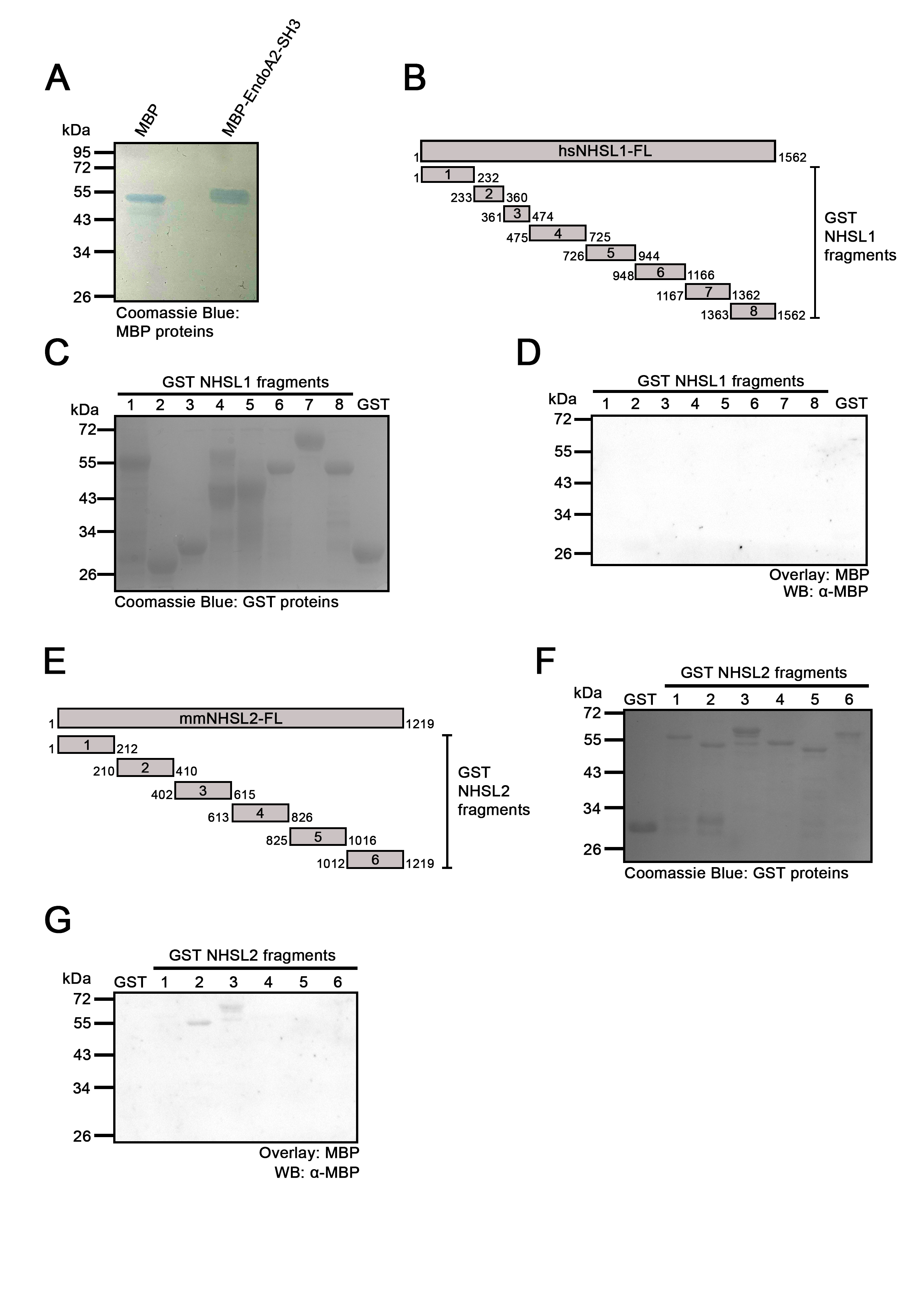

**Figure S2.** *(previous page)* **Related to Figure 3.** **Coomassie gels showing purity of MBP-EndoA2-SH3 and GST-tagged NHSL1 and NHSL2 fragments, and MBP-control far western blots.** (**A**) MBP-EndoA2-SH3 or MBP were purified from *E. coli* using amylose beads, separated via SDS-PAGE, and visualised with Coomassie blue stain. Note higher molecular weight of MBP-EndoA2-SH3 band compared to MBP, corresponding to additional 66 amino acids (~7kDa) from EndoA2-SH3.(**B**) Schematic showing 8 GST-fragments spanning the entire sequence of NHSL1 in relation to full length human NHSL1. The numbers refer to the amino acid position of the start and end of each fragment. (**C**) The 8 fragments were purified from *E. coli* using glutathione beads, separated via SDS-PAGE, and visualised with Coomassie blue stain. (**D**) Far western blot experiment using GST-tagged NHSL2 fragments overlaid with MBP control and probed using an antibody against MBP (exposed at the same time and with the same parameters as **Fig. 3A**). Blots are representative of three individual experiments. (**E**) Schematic showing 6 GST-fragments spanning the entire sequence of NHSL2 in relation to full length human NHSL2. The numbers refer to the amino acid position of the start and end of each fragment. (**F**) The 6 fragments were purified from *E. coli* using glutathione beads, separated via SDS-PAGE, and visualised with Coomassie blue stain. (**G**) Far western blot experiment using GST-tagged NHSL1 fragments overlaid with MBP control and probed using an antibody against MBP (exposed at the same time and with the same parameters as **Fig. 3D**). Blots are representative of three independent experiments.

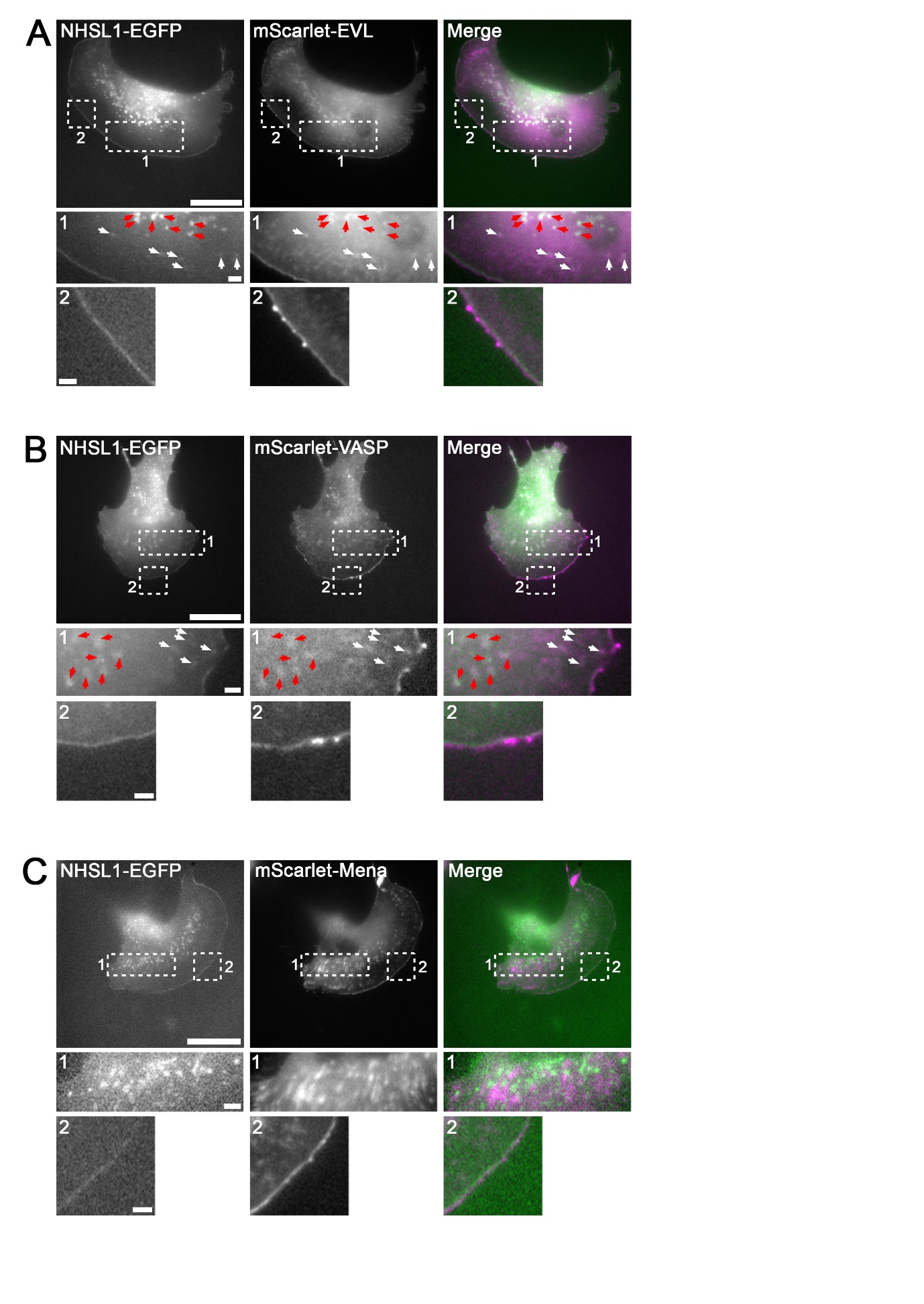

**Figure S3.** *(previous page)* **Related to Figure 4.** **NHSL1 co-localises with Ena/VASP proteins.** (**A** – **C**) B16-F1 cells were transfected with EGFP-tagged NHSL1 and mScarlet-tagged EVL (**A**, **Supplementary Movie 6**), VASP (**B**, **Supplementary Movie 7**) or Mena (**C**, **Supplementary Movie 8**) and plated on laminin. Localisation of tagged proteins was imaged live. White and red arrows mark distinct populations of vesicular puncta where co-localisation is observed. In merged stills, EGFP-tagged NHSL1 is represented in green, whilst mScarlet-tagged Ena/VASP proteins are represented in magenta. Scale bars: 20µm. Scale bar of insets: 2µm. Images representative of two (EVL, Mena) and three (VASP) independent experiments.

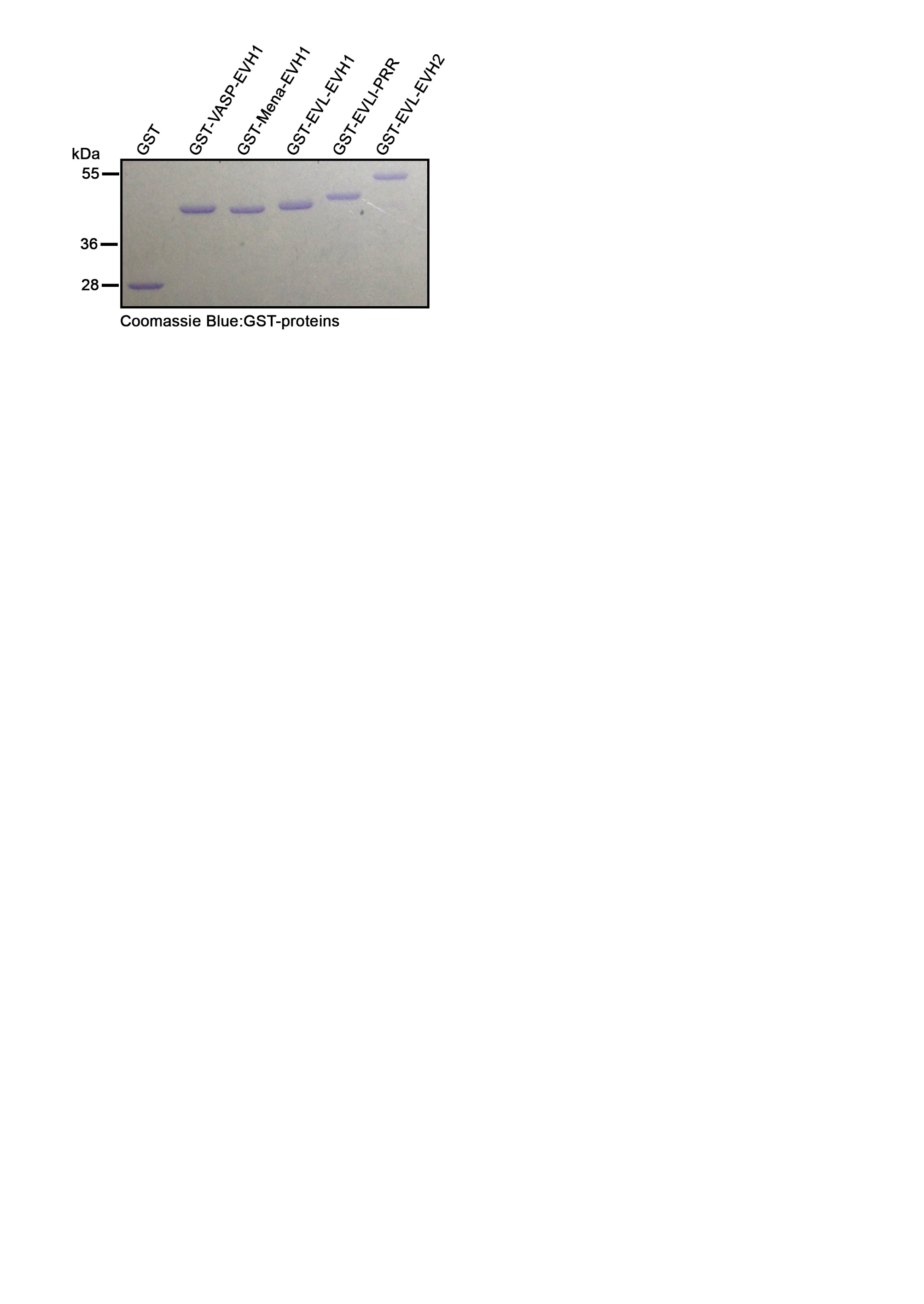

**Figure S4.** **Related to Figure 4. Coomassie gel showing purified subdomains of Ena/VASP proteins.** The EVH1 domains of EVL, VASP and Mena as well as the proline rich region (PRR) and EVH2 domain of EVL were purified from *E. coli* using glutathione beads, separated via SDS-PAGE and visualised with Coomassie blue stain.

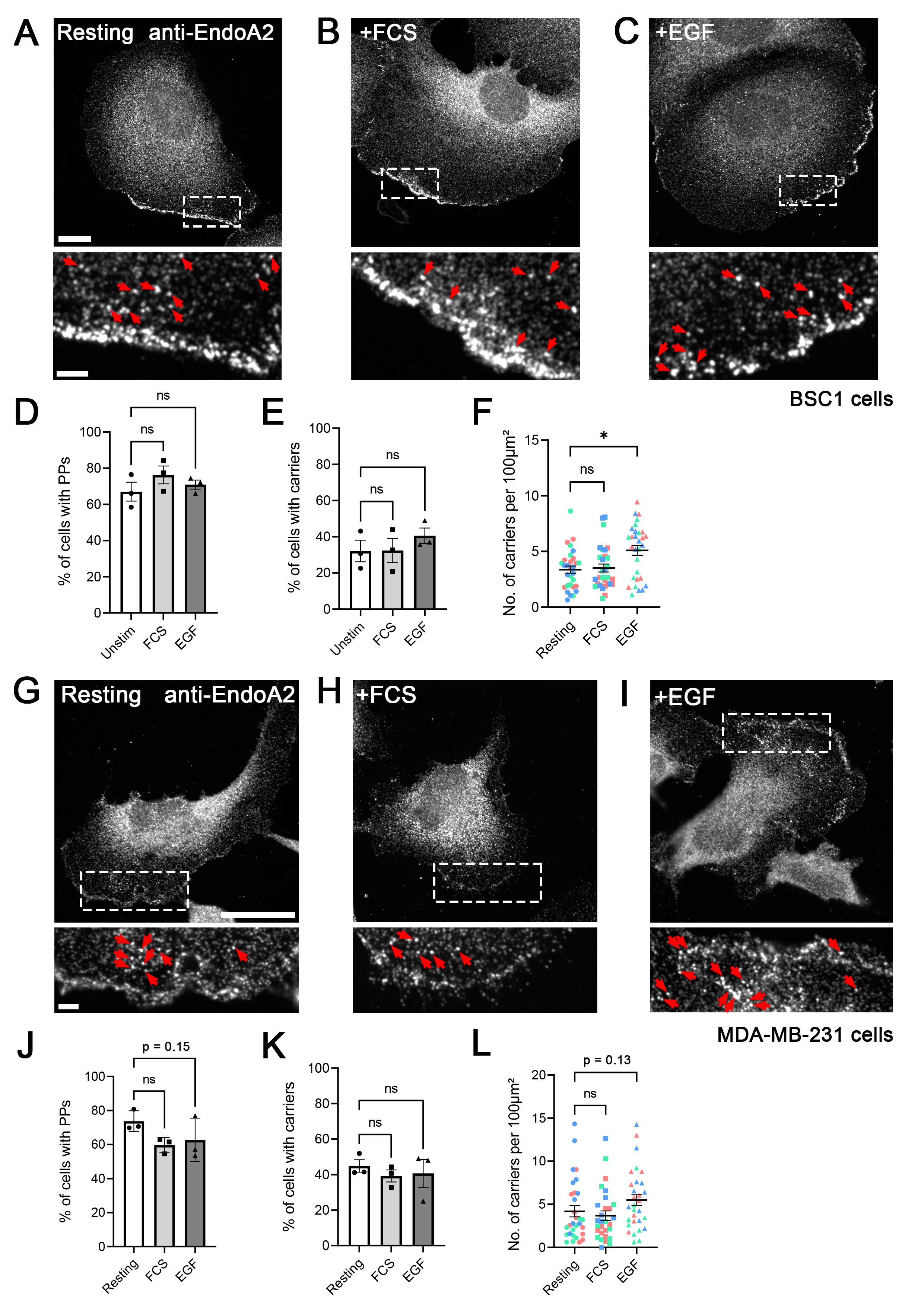

**Figure S5.** *(previous page)* **Quantification of FEME in BSC1 and MDA-MB-231 cells.** (**A** – **C**) BSC1 cells were grown in normal growth medium and then stimulated with media containing an additional 10% FCS (20% FCS total; **B**), 10ng/ml EGF (**C**) or left unstimulated (**A**). Cells were fixed, immuno-stained for EndoA2 and imaged at single confocal plane placed in the middle (in Z) of the lamellipodium. Scale bar: 10μm, Scale bar of inset: 2μm. Red arrows mark FEME carriers. (**D** – **E**) Graphs representing percentage of BSC1 cells containing FEME priming patches (PPs) (**D**) or FEME carriers (**E**). Data represents means ± SEM with percentages from individual biological replicates shown. N>150 cells per condition across 3 biological replicates. Statistics performed using one-way ANOVA with Dunnett’s multiple comparisons test, ns p>0.05. (**F**) Quantification of the number of FEME carriers per BSC1 cell normalised to a 100μm^2^ area. Data represents means + SEM with individual data points shown. Each data point represents 1 cell. Colours represent individual biological replicates. N = 30 cells per condition across 3 biological replicates. Statistics performed using Kruskal-Wallis test, *p=0.01, ns p>0.999. (**G** – **I**) MDA-MB-231 cells were grown in normal growth medium on collagen and then stimulated with media containing an additional 10% FCS (20% FCS total; **H**), 10ng/ml EGF (**I**) or left unstimulated (**G**). Cells were fixed, immuno-stained for EndoA2 and imaged at a single confocal plane placed in the middle (in Z) of the lamellipodium. Scale bar: 10μm, Scale bar of inset: 2μm. Red arrows mark FEME carriers. (**J** – **K**) Graphs representing percentage of MDA-MB-231 cells containing FEME priming patches (PPs) (**J**) or FEME carriers (**K**). Data represents means ± SEM with percentages from individual biological replicates shown. N>150 cells per condition across 3 biological replicates. Statistics performed using one-way ANOVA with Dunnett’s multiple comparisons test, ns p>0.05. (**L**) Quantification of the number of FEME carriers per MDA-MB-231 cell normalised to a 100μm^2^ area. Data represents means ± SEM with individual data points shown. Each data point represents 1 cell. Colours represent individual biological replicates. N = 30 cells per condition across 3 biological replicates. Statistics performed using Kruskal-Wallis test, ns p>0.05. P values under 0.2 are indicated on the figure.

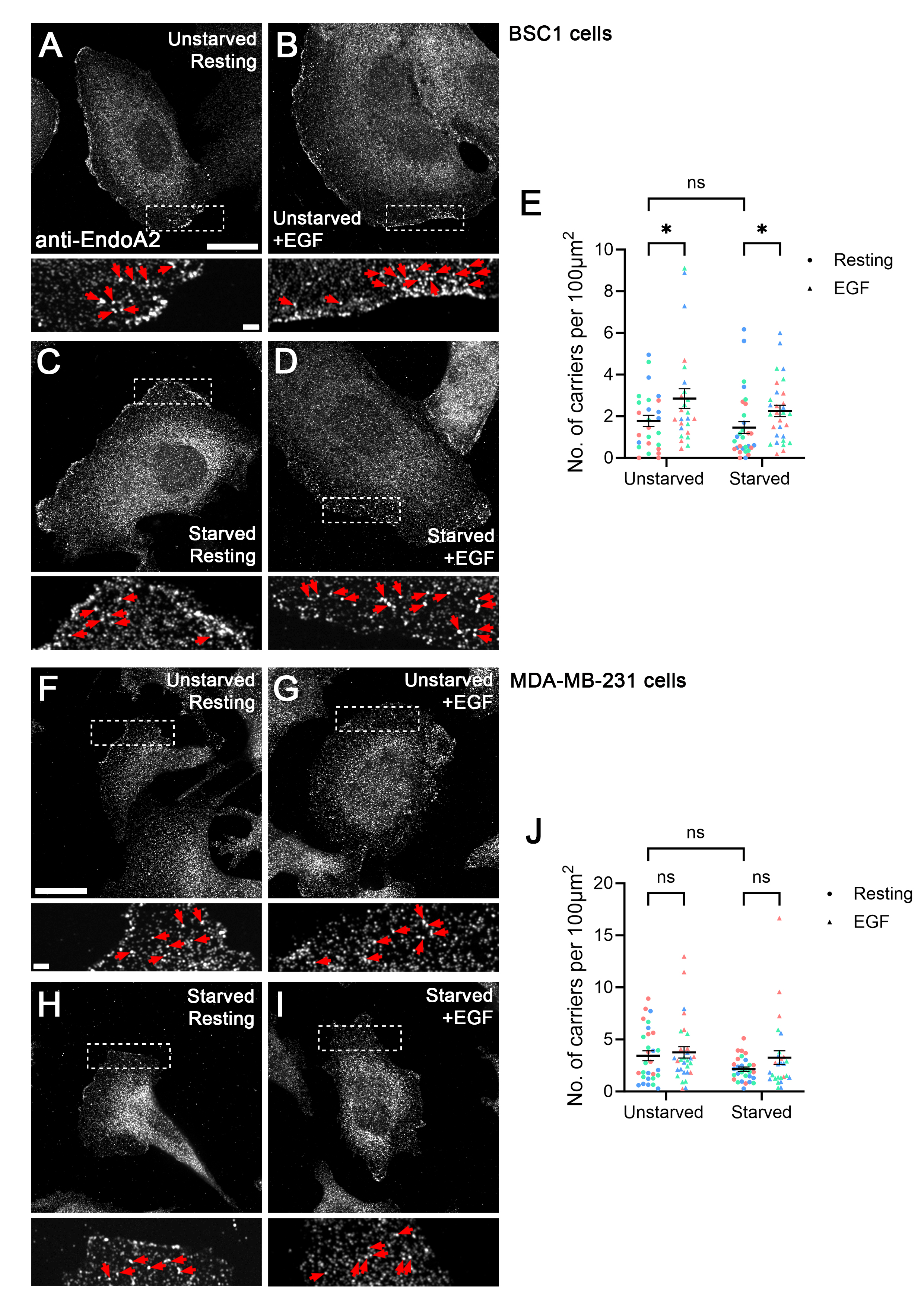

**Figure S6.** *(previous page)* **FEME response to short-term serum starvation in BSC1 and MDA-MB-231 cells** (**A** – **D**) BSC1 cells were initially grown in normal growth medium. Prior to fixation cells were incubated in normal growth medium with (unstarved, **A**, **B**) or without (serum starved, **C**, **D**) 10% FCS for 30 minutes and were subsequently stimulated with 10ng/ml EGF (**B**, **D**) or left unstimulated (**A**, **C**). Cells were fixed, immuno-stained for EndoA2 and imaged at a single confocal plane placed in the middle (in Z) of the lamellipodium. Scale bar: 20μm, Scale bar of inset: 2μm. Red arrows mark FEME carriers. (**E**) Quantification of the number of FEME carriers per BSC1 cell normalised to a 100μm^2^ area. Data represents means + SEM with individual data points shown. Each data point represents 1 cell. Colours represent individual biological replicates. N = 25-30 cells per condition across 3 biological replicates. Statistics performed using one-way ANOVA with Dunnett’s multiple comparisons test, *p<0.05. (**F** – **I**) MDA-MB-231 cells were initially grown in normal growth medium. Prior to fixation cells were incubated in normal growth medium with (unstarved, **F**, **G**) or without (serum starved, **H**, **I**) 10% FCS for 30 minutes and were subsequently stimulated with 10ng/ml EGF (**G**, **I**) or left unstimulated (**F**, **H**). Cells were fixed, immuno-stained for EndoA2 and imaged at a single confocal plane placed in the middle (in Z) of the lamellipodium. Scale bar: 20μm, Scale bar of inset: 2μm. Red arrows mark FEME carriers. (**J**) Quantification of the number of FEME carriers per MDA-MB-231 cell normalised to a 100μm^2^ area. Data represents means + SEM with individual data points shown. Each data point represents 1 cell. Colours represent individual biological replicates. N = 27-30 cells per condition across 3 biological replicates. Statistics performed using one-way ANOVA with Dunnett’s multiple comparisons test, ns p>0.05.

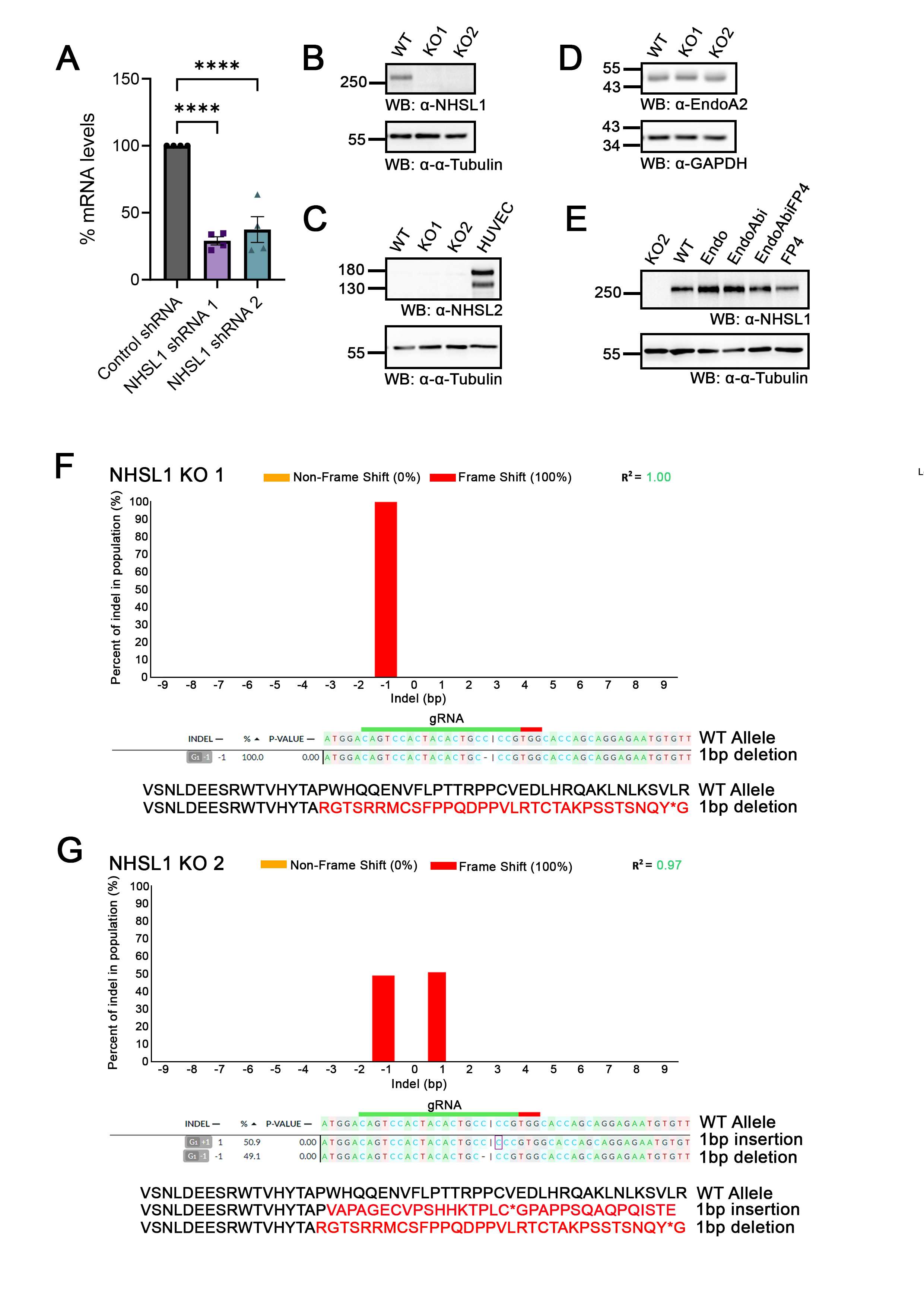

**Figure S7.** *(previous page)* **Related to Figure 6 & 7. Validation of NHSL1 knock-down and knock-out MDA-MB-231 cell lines and rescue constructs.** (**A**) mRNA levels of NHSL1 in MDA-MB-231 cells expressing shRNAs against NHSL1 or a control shRNA was assessed by qPCR using isoform independent, gene specific primer sets relative to expression of housekeeping gene B2M. Data represents means ± SEM with individual data points shown. N = 4 biological replicates. Statistics performed using one-way ANOVA with Dunnett’s multiple comparisons test: ****P < 0.0001. (**B** – **D**) Western blots showing protein levels of NHSL1 (**B**), NHSL2 (**C**, HUVEC lysates probed as a positive control for NHSL2 expression) and EndoA2 (**D**) in wild-type (WT), NHSL1 KO1 and KO2 MDA-MB-231 cells. Alpha-tubulin (α-tubulin) and GAPDH were used as loading controls. Blots are representative of three independent experiments. (**E**) Western blots showing NHSL1 protein levels in NHSL1 KO 2 MDA-MB-231 cells transfected with a Myc-only control plasmid (KO2) or the indicated NHSL1 constructs and selected with blasticidin. Alpha-tubulin (α-tubulin) was used as a loading control. Blots are representative of three independent experiments. (**F** – **G**) Genomic DNA was isolated from wild-type MDA-MB-231 cells as well as potential NHSL1 KO clones, and the region of DNA encompassing the expected cut site in exon 2 was amplified by PCR. These amplicons were sequenced, and DNA Sequences of knock-out clones were compared to WT sequence using the DECODR web tool. (**F**) 100% of alleles in NHSL1 KO 1 contain a deletion of one cytosine at the cut site resulting in a frame shift and a premature termination codon. (**G**) NHSL1 KO 2 contains one allele with a deletion of one cytosine, and one allele with an insertion of one cytosine at the cut site. Both mutations result in a frame shift and a premature termination codon. For both panels the entire amino acid sequence of exon 2 is shown in the wild-type mutant alleles. Altered amino acid sequence as a result of the frame shift is shown in red and the premature termination codon is represented with a ‘*’.

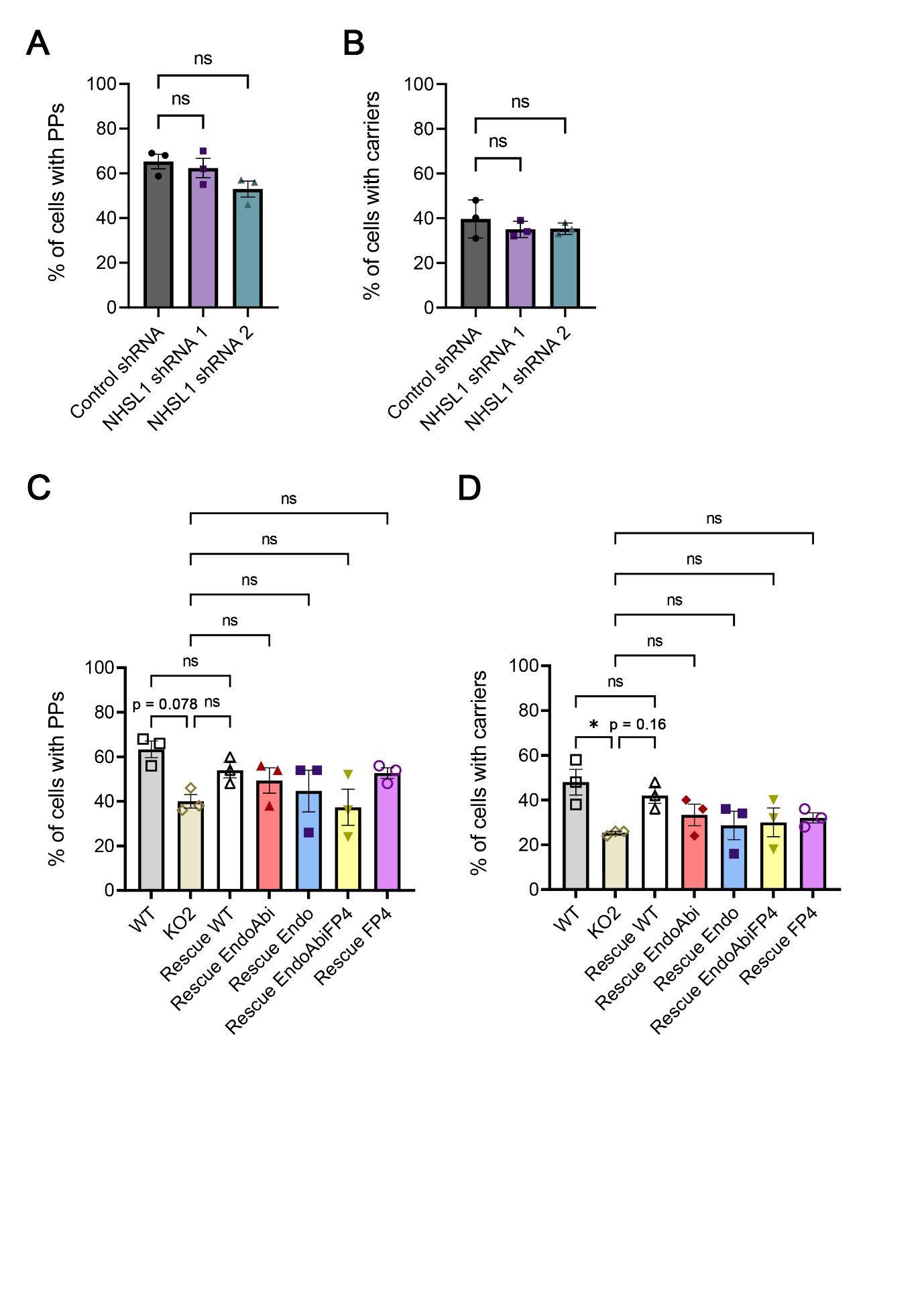

**Figure S8.** *(previous page)* **Related to Figure 6. Quantification of the percentage of NHSL1 knock-down and knock-out cells exhibiting FEME priming patches and carriers.** (**A** – **B**) Graphs representing the percentage of NHSL1 knock-down or control cells containing FEME priming patches (PPs) (**A**) or FEME carriers (**B**). Data represents means ± SEM with mean percentages from individual biological replicates shown. N=300 cells per condition across 3 biological replicates. Statistics performed using One-way ANOVA with Dunnett’s multiple comparisons test, ns p>0.05. (**C** – **D**) Graphs representing percentage of cells from knock-out-and-rescue experiments containing FEME priming patches (PPs) (**C**) or FEME carriers (**D**). Data represents means ± SEM with mean percentages from individual biological replicates shown. N=150 cells per condition across 3 biological replicates. Statistics performed using One-way ANOVA with Dunnett’s multiple comparisons test, ns p>0.05, *<0.05. P values under 0.2 are indicated on the figure.

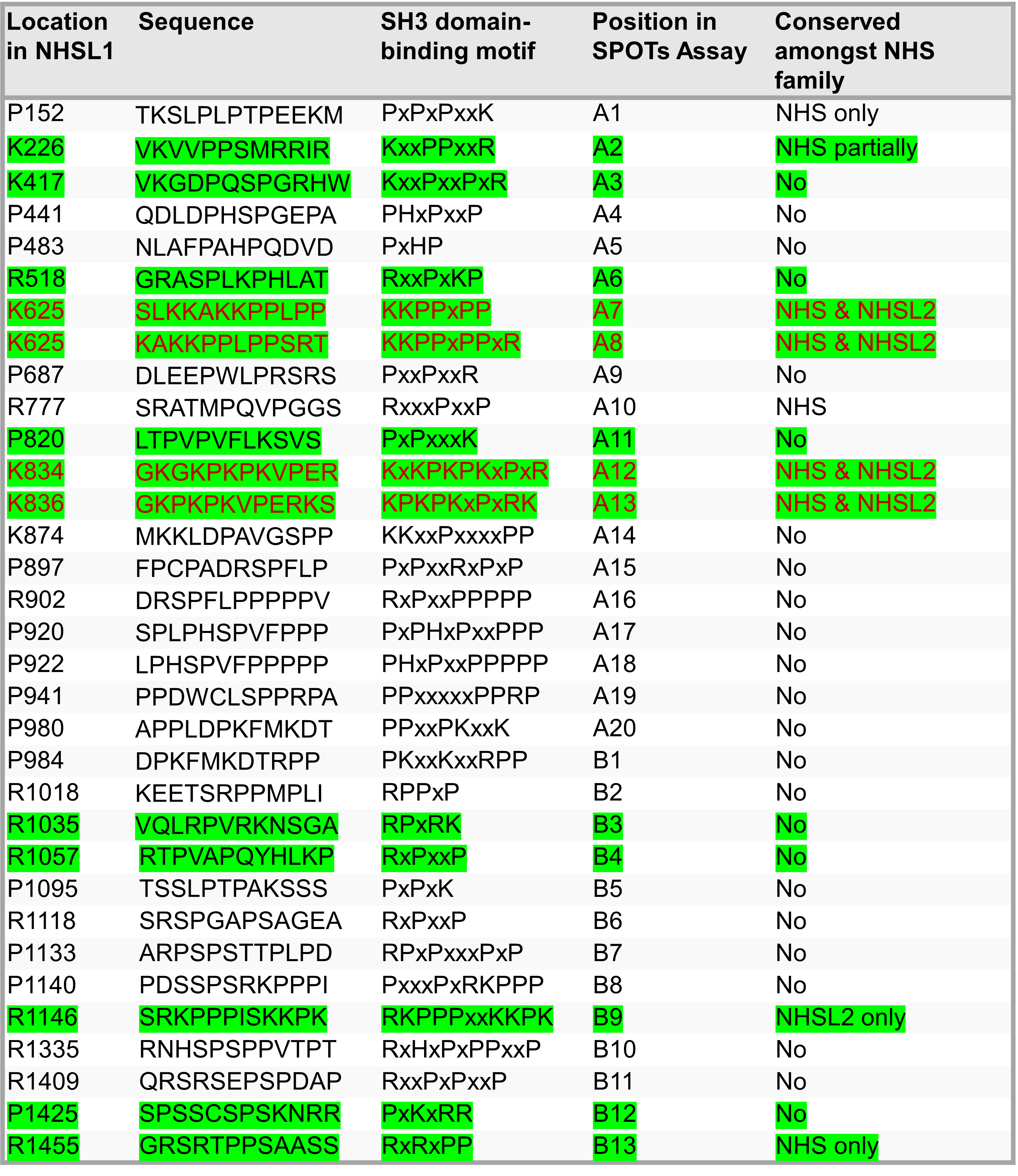

**Table S1. Putative SH3 domain-binding motifs in human NHSL1.** All 33 putative SH3 domain binding motifs in human NHSL1 identified using ScanSite. Sequence refers to 12mer peptide containing each motif assessed in peptide array (Fig. 3E). Sequence of the core motif is also indicated. Location refers to position of the first proline or positively charged residue of each motif relative to the entire length of human NHSL1. Conservation amongst NHS family was assessed using sequence alignment of NHSL1 with NHS and NHSL2. Positive hits in peptide array (Fig. 3E) are highlighted in green. Previously characterised Abi binding sites are shown in red.

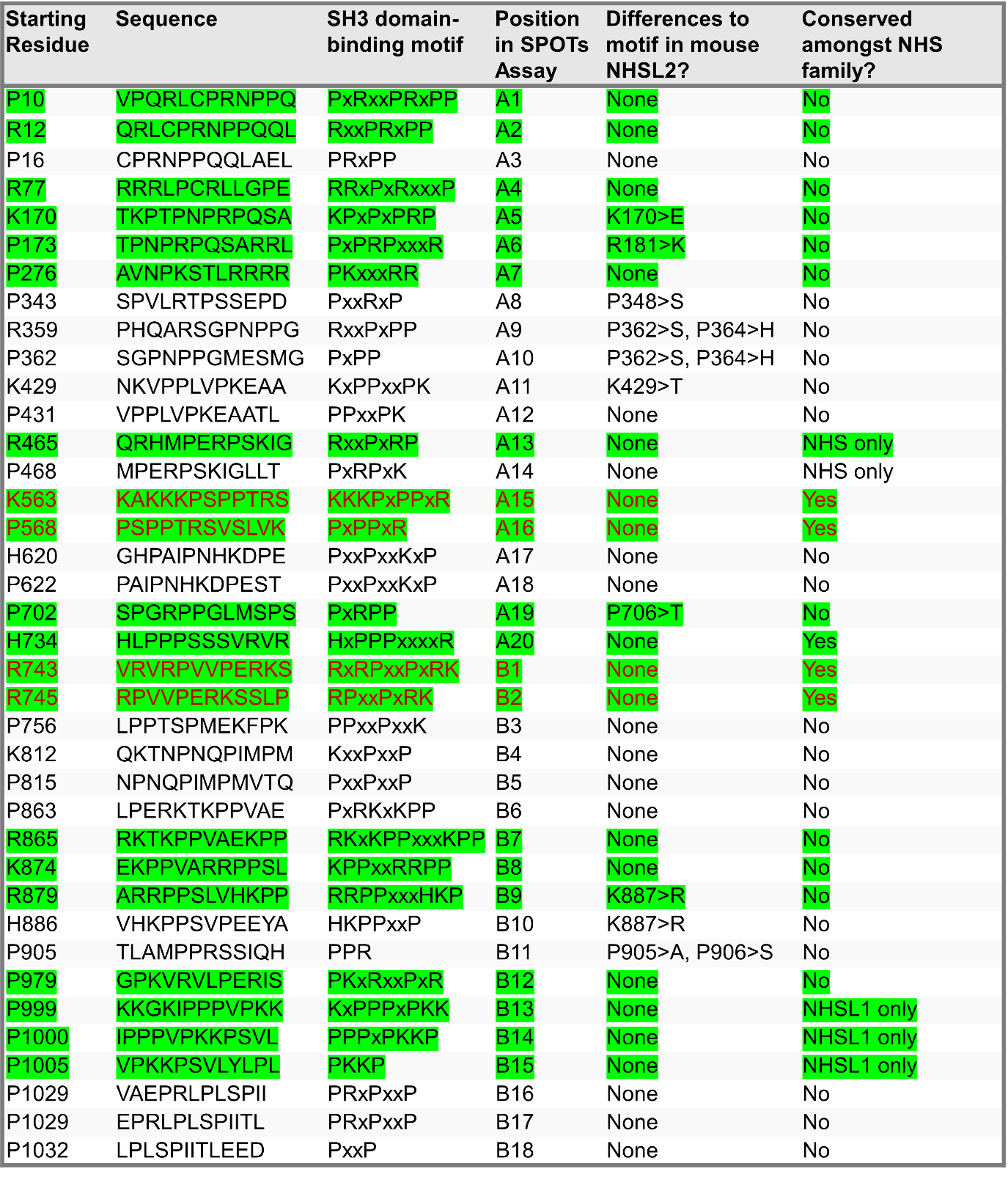

**Table S2. Putative SH3 domain-binding motifs in human NHSL2.** All 38 putative SH3 domain binding motifs in human NHSL2 identified using ScanSite. Sequence refers to 12mer peptide containing each motif assessed in peptide array (Fig. S1E). Sequence of the core motif is also indicated. Location refers to position of the first proline or positively charged residue of each motif relative to the entire length of human NHSL2. Differences in core motifs between human and mouse NHSL2 are specified for each site. Conservation amongst NHS family was assessed using sequence alignment of NHSL2 with NHS and NHSL1. Positive hits in peptide array (Fig. S1E) are highlighted in green. Previously characterised Abi binding sites are shown in red.
